# Cryo-EM structures of a neofunctionalized tardigrade peroxiredoxin specialized for nucleic acid binding

**DOI:** 10.64898/2026.04.10.717662

**Authors:** Haruka Yamato, Yohta Fukuda, Masanori Obana, Yasushi Fujio, Tsuyoshi Inoue

## Abstract

Some terrestrial tardigrades can endure severe oxidative stress in part through their gene-expanded repertoire of antioxidant proteins. However, among these antioxidant proteins, *Rv*PrxL, a peroxiredoxin (Prx)-like protein from *Ramazzottius varieornatus* strain YOKOZUNA-1, is unusual in that the catalytic cysteine is replaced by glutamate, apparently incapacitating canonical peroxidase function. In this study, we investigated the structure and function of this atypical Prx. Biochemical assays demonstrated that *Rv*PrxL completely lacks canonical peroxidase and chaperone activities. Cryo-EM analysis revealed a unique 20-mer structure of *Rv*PrxL. We also observed that the tardigrade-specific N-terminal region of *Rv*PrxL helps retain the high-order oligomeric assembly even in the presence of highly concentrated hydrogen peroxide. The N-terminal region also promotes nuclear localization of *Rv*PrxL and mediates binding to nucleic acids including nuclear RNAs. Furthermore, combining a standard cryo-EM method and a new approach in which cell **L**ysates are **Ap**plied **D**irectly **o**n cryo-EM **G**rids (LApDoG), we visualized two distinct nucleic acid–binding modes of *Rv*PrxL, termed “on-ring” and “in-ring”, which likely reflect distinct physiological roles or modes of action. Collectively, these findings show that *Rv*PrxL has been neofunctionalized to interact with nucleic acids in the nucleus, highlighting unexpected functional diversification of antioxidant proteins in tardigrades.

## Introduction

Tardigrades, also known as water bears, are microscopic animals that live in aquatic environments (1). They are known for their ability to survive under desiccated conditions by entering a unique anhydrobiosis (life without water) state, in which they lose their body water and stop metabolism (2, 3). In addition, anhydrobiotic tardigrades can withstand a variety of severe conditions, such as a wide range of temperatures, x-ray radiation, and high pressure (4). Therefore, anhydrobiotic tardigrades have been widely studied for their potential applications in both biomedical and industrial fields (5, 6). At the mechanistic level, accumulating evidence indicates that tardigrade proteins play central roles in establishing and maintaining anhydrobiosis. Consequently, extensive efforts have been devoted to identifying and characterizing tardigrade proteins that underpin anhydrobiosis and stress tolerance (7–11). Especially, their ability of radiation tolerance has attracted attention because it suggests the presence of unique unknown systems that protect or regulate nucleic acids (12–16).

Tardigrades go through severe oxidative stress under anhydrobiotic conditions (17); to cope with such stresses, tardigrades are thought to expand their repertoire of antioxidant proteins such as peroxiredoxin (Prx) and catalase (12, 18, 19), which decompose a reactive oxygen species (ROS), hydrogen peroxide (H_2_O_2_). In fact, the genome of *Ramazzottius varieornatus* strain YOKOZUNA-1, a model species of anhydrobiotic tardigrades and one of the toughest tardigrades, encodes nine putative Prx genes (12).

Prxs are enzymes that constitute a large family of peroxidase and play a major role in eliminating ROS in diverse organisms (20, 21). Prxs are classified into typical 2-Cys, atypical 2-Cys, and 1-Cys Prx subfamilies (22–24). Typical 2-Cys Prx can reduce peroxides such as H_2_O_2_ with conserved cysteine (Cys) residues through two reactions. First, its peroxidatic Cys (Cys^P^) reacts with peroxide, forming cysteine sulfenic acid (Cys^P^-SOH), and then, Cys^P^-SOH reacts with another Cys (resolving Cys or Cys^R^) to form a disulfide bond, which is eventually reduced by an appropriate electron donor (24–26). In addition to the role in scavenging ROS, a chaperone activity of typical 2-Cys Prxs such as cPrx1 from yeast, Prx1, Prx3 and Prx4 from *Homo sapiens* (*Hs*Prx1, *Hs*Prx3, and *Hs*Prx4, respectively) and Prx1 from *Schistosoma mansoni* (*Sm*Prx1) has been reported (27–31). These Prxs moonlight as a holdase which is one of the chaperone functions that prevents client proteins from denaturing and aggregating. 2-Cys Prx normally forms a ring structure composed of 10-mer or 12-mer to perform the antioxidative function, but when it goes through low pH or oxidative stress, it forms a filament-like higher molecular weight structure to function as a chaperone (31–33). Both ROS scavenging and chaperone functions indicate that 2-Cys Prxs are crucial for enduring extreme environments.

*Rv*PrxL, one of the putative Prxs from *R. varieornatus* YOKOZUNA-1, is predicted to have a flexible N-terminal region (amino acids 1–33) and a Prx-like region (amino acids 34–238) (Fig. 1A). The Prx-like region shows a high sequence identity with well-characterized 2-Cys Prxs (Supplementary Fig. S1); however, the deposited genomic and RNA-seq data show that *Rv*PrxL harbors a glutamate residue (Glu90) at the Cys^P^ position (Fig. 1B). We also independently confirmed this mutation by DNA sequencing of the *Rv*PrxL gene reverse-transcribed from mRNA purified from *R*. *varieornatus* (Supplementary Fig. S2A). Because Cys^P^ directly interacts with H_2_O_2_ and is hence indispensable for the catalytic reaction, the Cys-to-Glu substitution should abolish the enzymatic ability. Therefore, it is reasonable to ask this: does *Rv*PrxL perform the canonical role(s) of Prxs? If not, why has this gene been retained and evolved in *R*. *varieornatus* YOKOZUNA-1? This study sought to shed light on the function of *Rv*PrxL by combining structural analyses with functional characterization.

**Fig. 1.**
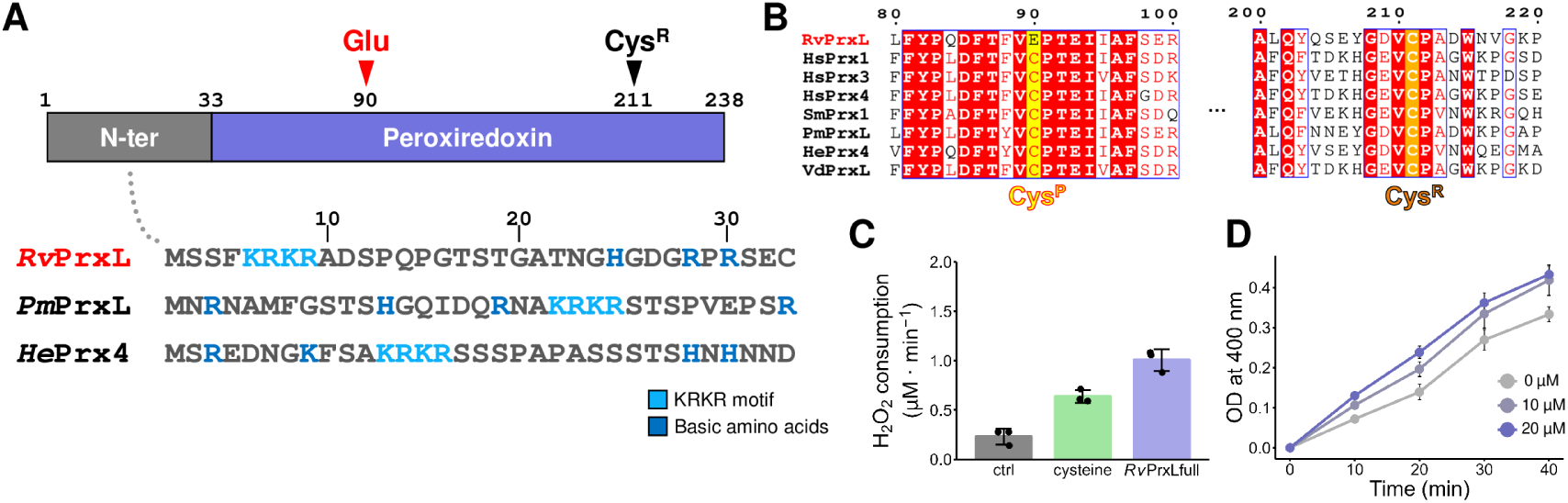
Canonical activity measurement of *Rv*PrxL. (**A**) Schematic of *Rv*PrxL and amino acid sequence of the N-terminal region (amino acids 1−33) of *Rv*PrxL, Prx2-like protein from *Paramacrobiotus metropolitanus* (*Pm*PrxL) and Prx4 from *Hypsibius exemplaris* (*He*Prx4). Blue alphabets are the basic amino acids: Lysine (K), Arginine (R), and Histidine (H). Cyan alphabets indicate the putative NLS motif (KRKR). (**B**) Comparison of the amino-acid sequence around “peroxidatic” Cys (Cys^P^) and “resolving” Cys (Cys^R^) to other homolog Prx-like proteins and well-characterized typical 2-Cys Prxs; Prx1, Prx3, Prx4 from *H. sapiens* (*Hs*Prx1, *Hs*Prx3, *Hs*Prx4, respectively), Prx1 from *S. mansoni* (*Sm*Prx1), Prx2-like protein from *P. metropolitanus* (*Pm*PrxL), Prx4 from *H. exemplaris* (*He*Prx4), and peroxiredoxin-like isoform X1 from *Varroa destructor* (*Vd*PrxL). Regarding *Rv*PrxL, Glu is encoded on the position of Cys^P^ of other Prxs. (**C**) H_2_O_2_ consumption per minute of control, L-cysteine, and *Rv*PrxL, measured using the FOX assay method. 100 µM dithiothreitol was used as a reductant to complete the catalytic cycle of Prx. Assay data are mean ± SD of three technical replicates for each assay. (**D**) Results of the holdase activity assay by tracking the 400 nm absorbance of reduced lysozyme. 0 µM (gray), 10 µM (light purple), 20 µM (purple) concentration of *Rv*PrxL is added to the reaction mixture, respectively. Assay data are mean ± SD of three technical replicates for each assay.

## Materials and Methods

### Sequence analysis

Protein sequences of interest were obtained from The National Center for Biotechnology Information (NCBI) database (https://www.ncbi.nlm.nih.gov/). BLAST searches were conducted using the NCBI protein BLAST web service (https://blast.ncbi.nlm.nih.gov/Blast.cgi) (34). Sequence alignments were performed with Clustal Omega (https://www.ebi.ac.uk/jdispatcher/msa/clustalo). Visual representations of the sequence alignments were generated using ESPript 3.0 (http://espript.ibcp.fr) (35).

Phylogenetic analysis of *Rv*PrxL was performed using MEGA X (36). A maximum likelihood method was used to estimate a phylogenetic tree, incorporating 500 bootstrap replicates using the Jones-Taylor-Thornton model. A figure of the phylogenetic tree was illustrated by using iTOL v6 (https://itol.embl.de/) (37). The list of the species for the phylogenetic analysis and GenBank accession numbers are provided in Supplementary Table S1.

Nuclear localization signal (NLS) sequences were predicted using NLS Mapper (https://nls-mapper.iab.keio.ac.jp/) (38) and NLSExplorer (http://www.csbio.sjtu.edu.cn/bioinf/NLSExplorer/index.html) (39). The results are shown in Supplementary Table S2.

## Cloning, expression, and purification of *Rv*PrxL

*R. varieornatus* strain YOKOZUNA-1 was generously provided by T. Kunieda at The University of Tokyo (currently at University of Hyogo). The expression plasmids for full-length *Rv*PrxL (Gene name: RvY_14602, Genbank ID: GAV04299.1) and ΔN_1–33_ mutant proteins were prepared using an identical protocol. A complementary DNA (cDNA) library was prepared as previously described (11). The *Rv*PrxL genes were amplified from cDNA, subsequently cloned into the pET-28a(+) vector (Novagen) via the In-Fusion cloning method (Takara Bio). The obtained plasmid vector was amplified in *Escherichia coli* strain DH5α and purified with a NucleoSpin Plasmid EasyPure kit (MACHEREY-NAGEL). Primers used for PCR are noted on Supplementary Table S3. All DNA constructs were verified by DNA sequencing (Fasmac Co., Ltd.). The pET-28a(+)–*Rv*PrxL plasmids were transformed into *E. coli* expression strain BL21(DE3) cells and cultured in LB medium supplemented with 100 µg/mL kanamycin sulfate (FUJIFILM Wako Pure Chemical Corporation) at 37 °C until OD_600_ reached 0.6. The protein expression was induced by adding 0.1 mM isopropyl β-D-1-thiogalacto pyranoside (IPTG) (Nacalai Tesque), followed by incubation for 20 hours at 18 °C. Cells were collected by centrifugation (6, 000×g, 30 min) at 4 °C and resuspended in Buffer A (20 mM Tris-HCl pH 8.0, 200 mM NaCl and 10% glycerol). Cell lysis was performed by sonication on ice, and the lysate was centrifuged (20, 000×g, 60 min) at 4 °C to remove cellular debris.

The proteins were initially intended to be purified by incorporating a 6×His-tag and a TEV protease cleavage site. However, removal of the His-tag employing TEV protease was unsuccessful, regardless of whether the tag was positioned at the N- or C-terminus. This failure may be attributed to the protein’s high oligomeric state, potentially resulting from interactions within the N-terminal and C-terminal regions with other *Rv*PrxL molecules. Consequently, untagged proteins were purified using the following purification protocol. The obtained sample was first subjected to ammonium sulfate precipitation. Prior to the main experiment, a small aliquot of the sample was used to determine the ammonium sulfate concentration at which *Rv*PrxL precipitates by performing stepwise precipitation with increasing ammonium sulfate concentrations. First, the supernatant was treated with 30% ammonium sulfate and centrifuged (20, 000×g, 15 min) at 4 °C. Then, the supernatant was treated with 50% ammonium sulfate and centrifuged under the same condition.

The precipitate was resuspended and applied to a HiTrap Phenyl HP column (Cytiva) pre-equilibrated with Buffer B (20 mM Tris-HCl pH 8.0, 200 mM NaCl, 10% glycerol and 30% ammonium sulfate). The column was subsequently washed with Buffer B and eluted with Buffer A. The eluted fraction was collected and dialyzed overnight at 4 °C against Buffer C (20 mM Tris-HCl, pH 8.0 and 10% glycerol). The sample was then loaded onto Hitrap Q HP column (Cytiva) pre-equilibrated with Buffer C. The column was washed with Buffer C and proteins were eluted with a linear gradient from Buffer C to Buffer D (20 mM Tris-HCl pH 8.0, 1 M NaCl and 10% glycerol). The eluted fraction containing the target protein was then loaded onto a HiLoad 16/60 Superdex 200 prep grade column (GE Healthcare), which had been pre-equilibrated with Buffer E (20 mM Tris-HCl pH 8.0 and 200 mM NaCl) and eluted with the same buffer (Supplementary Fig. S2B). The target fraction was collected, concentrated, and stored at –80 °C.

Ultraviolet-visible (UV–vis) absorption spectra were recorded at each purification step using a NanoDrop One spectrophotometer (Thermo Fisher Scientific). The target fraction was assessed at each elution step by SDS-PAGE. For Clear Native PAGE (CN-PAGE), each protein sample was diluted to a concentration of 0.5 mg/mL.

### Sample preparation for the Lysates are Applied Directly on cryo-EM Grids (LApDoG) method

Full-length *Rv*PrxL was expressed in *E. coli* BL21(DE3) as described above. Cell lysis was performed by sonication on ice, and the lysate was centrifuged (20, 000×g, 60 min) at 4 °C to remove cellular debris. After that, the supernatant was filtered using a Minisart (0.45 μm / Luer Slip) Syringe filter (Sartorius) and dialyzed in Buffer E overnight. 3 µL of the obtained sample (Supplementary Fig. S2C) was loaded on a cryo-EM grid as described in the next section. The schematic illustration of the LApDoG method is shown in Supplementary Fig. S2D.

## Cryo-EM specimen preparation, data collection, and image processing

The purified *Rv*PrxL was diluted to 0.5 mg/mL in Buffer E, except for the RNA complexes, for which we used 0.3 mg/mL *Rv*PrxL and 1 mg/mL of either tRNA (Thermo Fisher Scientific) or U3/U6 RNA (see “Mobility shift assay and nuclease protection assay” section below for preparation details). For the H_2_O_2_ exposed structures, H_2_O_2_ was added as a final concentration of 100 mM, and the sample was incubated for 30 min at room temperature. All samples were prepared in Buffer E.

QUANTIFOIL R1.2/1.3 Cu 200 Grids (Quantifoil Micro Tools GmbH) were freshly glow-discharged by a JEC-3000FC Auto Fine Coater (JEOL). For observing the ΔN_1–33_ mutant treated with 100 mM H_2_O_2_, UltrAuFoil R1.2/1.3 Grid (Quantifoil Micro Tools GmbH) was used instead. Vitrobot Mark IV (Thermo Fisher Scientific) was used for blotting and cryocooling the grids. The chamber was kept at 4 °C and 100% humidity. 3 µL of the solution was applied onto the grid, incubated for 30 seconds in the chamber and blotted for 3 seconds before plunging into liquid ethane. Cryo-EM images of samples other than the U6 complex were collected using SerialEM (40) on CRYO ARM™ 200 (JEOL) electron microscope operated at 200 kV equipped with a K3 direct electron detector (Gatan) at the calibrated pixel size of 0.83 Å. Cryo-EM images of the U6 complex sample were collected using SerialEM on CRYO ARM™ 300 (JEOL) electron microscope operated at 300 kV equipped with a K3 direct electron detector (Gatan) at the calibrated pixel size of 0.86 Å. Data processing was performed in CryoSPARC (41). The detailed processing workflows and the number of micrographs used for analyses are noted on Supplementary Figs. S3−8 and Supplementary Tables S4−6.

## Model Building

An initial model for model building of full-length *Rv*PrxL was predicted by Colabfold (42). Full-length *Rv*PrxL (PDB ID: 23PU) was used for model building of *Rv*PrxL treated with H_2_O_2_ and the ΔN_1–33_mutant. Each initial models were fitted to the EM density maps using “fit in map” in UCSF Chimera (43) or UCSF ChimeraX (44). After that, each model was refined by Phenix (45, 46) and manually adjusted using Coot (47). Then the models were further refined by Phenix and Servalcat (48). This round of refinement was conducted several times.

The structure of a zinc-finger domain containing Prx from *Hypsibius exemplaris* (Genbank ID: OQV14346.1) was predicted by Alphafold3 by using default parameters (49).

All structural figures are visualized and generated by UCSF ChimeraX or PyMOL (https://www.pymol.org/). The statistics for all data collection and structure refinement are summarized in Supplementary Tables S4−6.

## Peroxidase activity assay

All absorbance measurements were conducted using a SpectraMax M2 UV-Visible spectrophotometer operated with SoftMax Pro software (Molecular Devices).

Antioxidant activity was assessed by measuring OD at 240 nm (OD_240_) of H_2_O_2_ and by using the FOX assay method (50). OD_240_ values were tracked to evaluate antioxidative activity at higher concentrations of H_2_O_2_. Conversely, the FOX assay was employed to quantify the consumption of H_2_O_2_ at lower concentrations (<200 µM) with high sensitivity.

Firstly, we compared the antioxidant activities of *Rv*PrxL and bovine liver catalase (FUJIFILM Wako Pure Chemical Corporation) by measuring OD_240_. The reaction mixtures were prepared for each protein using the same protocol as the FOX assay. Immediately after the addition of 10 mM H_2_O_2_ to the mixtures, OD_240_ was recorded by a kinetic mode using a silica quartz square cell cuvette maintained at 37 °C.

For the FOX assay, FOX reagent A (25 mM ammonium ferrous sulfate and 2.5 M sulfuric acid) and FOX reagent B (50 mM sorbitol and 125 µM xylenol orange) were freshly prepared prior to the experiment. The reaction mixture was assembled in Buffer E within microfuge tubes, containing 5 µM of *Rv*PrxL or free L-cysteine (FUJIFILM Wako Pure Chemical Corporation), and 100 µM dithiothreitol (DTT; FUJIFILM Wako Pure Chemical Corporation), as DTT is essential for completing the Prx catalytic cycle. The control sample also contained 100 µM DTT. The tubes were incubated at 37 °C for 15 min immediately after adding 100 µM H_2_O_2_. Subsequently, 20 µL of the reaction mixture was added to 200 µL FOX working reagent (1:100 volume ratio mixture of FOX reagent A and FOX reagent B) to quench the reaction. The samples were vortexed and incubated at room temperature for 1h to stabilize. Absorbance at 560 nm was measured and H_2_O_2_ consumption for each sample was scaled using a standard curve generated from six different H_2_O_2_ concentrations.

## Holdase activity assay

All absorbance measurements were conducted using a SpectraMax M2 UV-Visible spectrophotometer operated with SoftMax Pro software. Holdase activity was assessed using the Lysozyme Aggregation Assay (51). First, the aggregation of 10 µM lysozyme (from egg white; FUJIFILM Wako Pure Chemical Corporation) was induced by the addition of 1 mM Tris(2-carboxyethyl)phosphine hydrochloride (Hampton Research) in Buffer E. OD at 400 nm (OD_400_) was monitored in a kinetic mode to quantify lysozyme aggregation. To evaluate holdase activity, either 10 µM or 20 µM of *Rv*PrxL was added prior to the reduction step. For assay control, 20 µM of Bovine serum albumin (Takara Bio) was used instead of *Rv*PrxL.

## Subcellular localization

The genes of full-length *Rv*PrxL, the ΔN_1–33_ mutant, and the KRKR-4A mutant were cloned into a pAcGFP-N1 vector (Clontech). The experiment was performed using the protocol based on our previous research (11). In the experiment, 3000 ng/well of *Rv*PrxL–pAcGFP-N1, ΔN_1–33_ mutant– pAcGFP-N1 and KRKR-4A–pAcGFP-N1 were transfected respectively into human embryonic kidney cells 293 (HEK293, RRID: CVCL_0045), obtained from RIKEN BRC (Tsukuba, Ibaraki, Japan).

## Mobility shift assay and nuclease protection assay

The synthesized genes (gBlocks Gene Fragment) of U3 and U6 RNA sequences were purchased from Integrated DNA Technologies (Coralville, IA, USA). The gene construct was starting with 8 nucleotides (CGCGAAAT) to induce the binding of T7 polymerase, followed by T7 binding nucleotides (TAATACGACTCACTATAGGG), and the target sequences which are described in Supplementary Table S7. RNA in vitro transcription was performed using an in vitro Transcription T7 Kit (Takara Bio) according to the user manual. The length of the RNAs were confirmed by the agarose gel electrophoresis.

Double-stranded DNA (dsDNA) was prepared by annealing complementary single-stranded DNAs (ssDNAs). The same amount of complementary ssDNAs were annealed by stepwise cooling from 95 °C to 25 °C over 1 h.

*Rv*PrxL and the ΔN_1–33_ mutant were incubated in Buffer E (20 mM Tris-HCl pH 8.0 and 200 mM NaCl) for 10 min on ice with the following amount of nucleic acids: 100 ng of plasmid (pET-28a(+), Novagen), 1 µg of ssDNA and dsDNA, 1 µg tRNA from yeast (Thermo Fisher Scientific) and 1 µg of U3 RNA and U6 RNA. The samples were separated with the agarose gel electrophoresis method using a 1.0% agarose gel (Lonza) and then stained with SYBR Gold (Thermo Fisher Scientific) to visualize the shift of nucleic acids. Nucleic acid sequences are described in Supplementary Table S7.

For nuclease protection assay, 5 U of recombinant DNase I (Takara Bio) was added to each reaction mixture with Buffer F (20 mM Tris-HCl pH 8.0, 200 mM NaCl and 8 mM MgCl_2_). The mixture was incubated at room temperature for 15 min. The intensity of each band was quantified using ImageJ (52). The remaining dsDNA ratio was calculated as the existing intensity relative to that of the control group.

## Results

### An impaired peroxiredoxin from *R. varieornatus*

Activity assays were performed to determine whether *Rv*PrxL functions as a typical Prx or as a chaperone. First, we measured changes of OD_240_ to monitor decomposition of H_2_O_2_ and found that the H_2_O_2_ consumption of *Rv*PrxL was approximately 10³-fold lower than that of catalase (Supplementary Fig. S9A). We also measured the consumption of H_2_O_2_ by the FOX assay. *Rv*PrxL only shows approximately 2 times more consumption of H_2_O_2_ than free L-cysteine (Fig. 1C), suggesting that H_2_O_2_ consumption by *Rv*PrxL derived from the oxidation of two cysteine residues existing in its polypeptide chain, rather than performing the enzymatic reaction.

Next, we performed a chaperone activity assay. We monitored OD_400_ to observe aggregation of lysozyme induced by disulfide bond reduction. When bovine serum albumin with known holdase function (53) was employed as an assay control, an inhibition of the increase in OD_400_ was observed (Supplementary Fig. S9B). However, addition of *Rv*PrxL in the reaction mixture did not reduce the increase of OD_400_, showing that *Rv*PrxL does not have the holdase function (Fig. 1D).

These results indicate that *Rv*PrxL completely lacks canonical Prx functions. At first glance, this appears inconsistent with the idea that anhydrobiotic tardigrades expand antioxidant enzymes to cope with oxidative stress. However, our findings instead suggest that the expanded repertoire of Prx-like proteins in tardigrades may include members that have diverged from canonical Prx functions and acquired distinct roles. To gain insight into the function of *Rv*PrxL, we next pursued structural analysis.

## Cryo-EM structural analyses of *Rv*PrxL

When we observed full-length *Rv*PrxL with cryo-EM, we found that some molecules formed short filament-like structures composed of stacked rings. However, we focused on a most abundant double-ring stacked state and determined its structure at 2.65 Å resolution as a representative architecture (Supplementary Fig. S3 and Table S4). This major structure of full-length *Rv*PrxL shows a 20-mer assembly, which is formed by the stacking of two 10-mer rings (Fig. 2A and Supplementary Fig. S9C). Whereas other homologous Prxs form a stacking structure only at low pH or under high concentration of H_2_O_2_ (31–33), *Rv*PrxL tends to stack even at pH 8.0 and in ambient conditions. Previous studies on Prxs have used mutations substituting Cys^P^ with a carboxyl-group-containing residue to mimic the hyperoxidized state of Cys^P^ (Cys^P^-S-OOH) (54). Therefore, Glu90 in *Rv*PrxL may naturally mimic the hyperoxidized state. Considering that Prxs are known to undergo higher-order oligomerization under oxidative stress, the tendency of *Rv*PrxL to form stacked structures under native conditions may be attributable to this residue substitution. In fact, the Cys^P^-to-Asp mutation in *Sm*Prx1 is known to enhance the stacking of 10-mer rings (55).

**Fig. 2.**
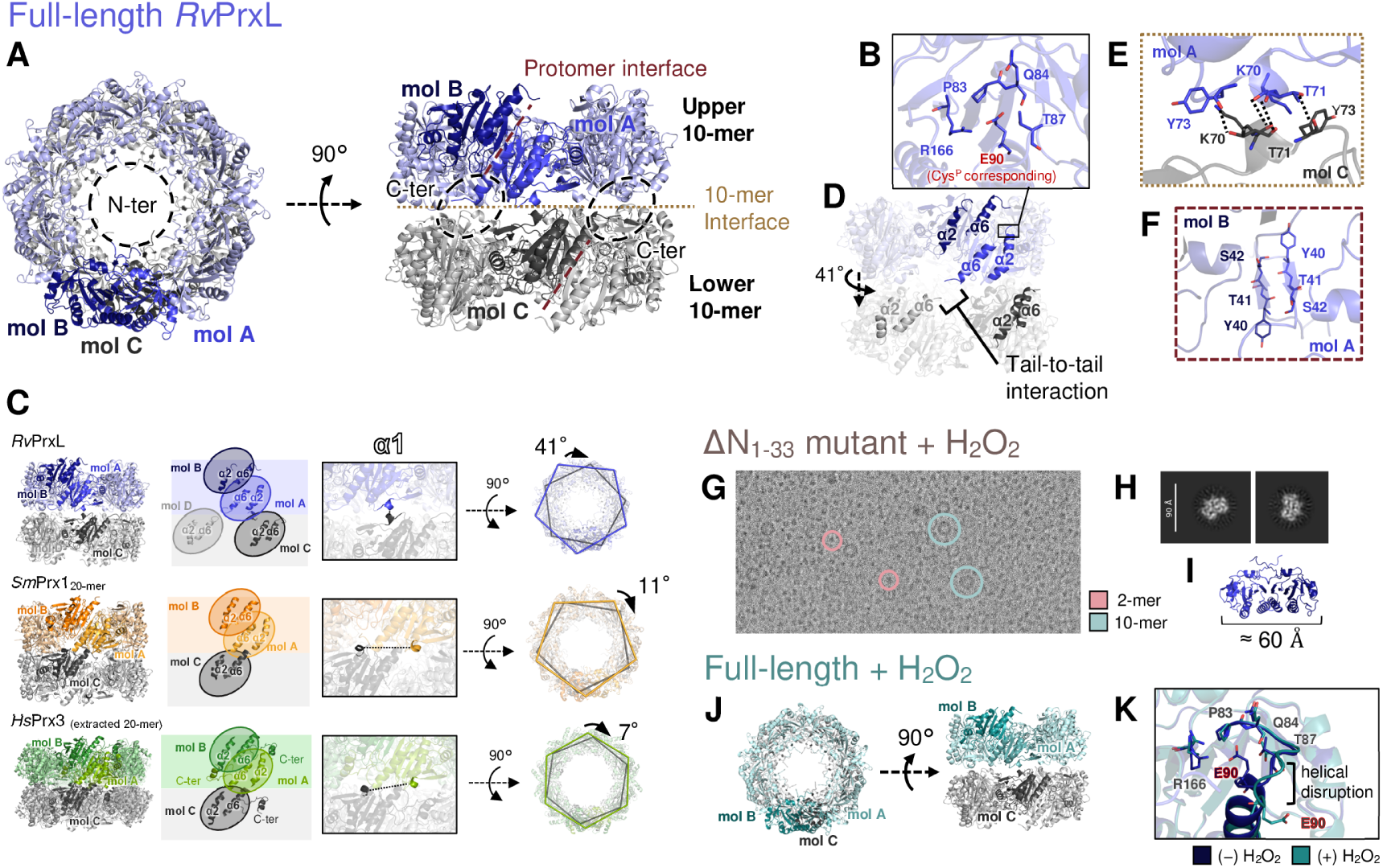
Cryo-EM structures of *Rv*PrxL. (**A**) Ribbon representation of full-length *Rv*PrxL, viewed from the top and after a 90° rotation about the horizontal axis. Molecules A, B, and C (mol A/B/C) are shown in darker color to highlight the protomer arrangement. (**B**) Glu90 corresponding to Cys^P^ and its adjacent residues. Oxygen and nitrogen atoms are colored red and blue, respectively. (**C**) Comparison of stacking manners observed in *Rv*PrxL, *Sm*Prx1, and *Hs*Prx3 (20-mer extracted from a filament structure). From left to right: overall structure, schematic of arrangement of α2 and α6 helices, separation distances between α1 helices, and rotation of the upper ring relative to the lower ring. To highlight the stacking arrangement, Cα atoms of Ala102, Ser64 and Val64 of *Rv*PrxL, *Sm*Prx1, and *Hs*Prx3 are joined respectively by dashed lines. (**D**) Intermolecular interaction between mol A and mol C at the interface between 10-mers. (**E**) Possible hydrogen bonds between mol A and mol B. (**F**) Molecular interface between mol A and mol B that forms a short β-sheet structure. (**G**) A representative micrograph of the ΔN_1–33_ mutant treated with H_2_O_2_. (**H**) The representatives of 2D classification images of small molecules of the ΔN_1–33_ mutant treated with 100 mM H_2_O_2_. (**I**) A 2-mer model in the full-length *Rv*PrxL structure. (**J**) Overall structure of full-length *Rv*PrxL treated with 100 mM H_2_O_2_. (**K**) Structural comparison between structures with and without exposure to H_2_O_2_. A minor loop rearrangement was induced by H_2_O_2_ treatment.

The monomeric secondary structure shows high structure similarity to known Prxs such as *Sm*Prx1 (PDB ID: 3ZVJ) and *Hs*Prx3 (PDB ID: 5JCG) with RMSD values of 0.8 and 0.6 Å, respectively. Glu90, the Cys^P^ corresponding residue, is located at the same position as Cys^P^ observed in other 2-Cys Prxs (Fig. 2B and Supplementary Fig. S9D).

Notably, the 20-mer assembly of *Rv*PrxL exhibits a novel stacking mode distinct from those observed in the previously reported Prxs. At the stacking interface between 10-mers of known Prxs, there is an interaction between α2 and α6, forming two conjugated long helices (Fig. 2C). Instead, in *Rv*PrxL, α6 from the adjacent protomers approach each other, forming tail-to-tail interaction via the subsequent C-terminal disordered regions (Figs. 2C−D, and Supplementary Fig. S9E). This arrangement in the *Rv*PrxL 20-mer results in a 41 degrees rotation of the upper 10-mer around the five-fold axis relative to the lower 10-mer, representing a distinct stacking geometry in the Prx family (Fig. 2C). This stacking mode of *Rv*PrxL places the α1 helices from adjacent rings in close proximity. Several residues from each protomer in this region are positioned within ∼3 Å of one another, suggesting potential hydrogen-bonding interactions (Fig. 2E). Although α1 is also present in *Hs*Prx3 and *Sm*Prx1, the α1 helices at the corresponding inter-ring interface are far apart in these proteins (Fig. 2C). In addition, *Rv*PrxL exhibits a unique interaction on the inner side of the ring, where a short β-sheet (β1) forms between neighboring protomers (Fig. 2F). The regions further toward the N-terminus are disordered and appear to interact with one another within the ring interior (Supplementary Fig. S9E).

To further investigate the structural feature of the N-terminal region, an N-terminal deletion (ΔN_1–33_) mutant was analyzed by cryo-EM (Supplementary Figs. S4, S10A−B and Table S4). Although the ΔN_1–33_ mutant maintains the same stacking mode of rings, some 10-mer rings assemble into extended filaments in addition to forming 20-mers. As a result, we obtained a map for the filament structure at 3.02 Å resolution. This tendency of the ΔN_1–33_ mutant to form filaments is likely due to altered electrostatic property and the loss of electrostatic repulsion between the positively charged N-terminal regions.

Exposure to 100 mM H_2_O_2_ revealed that the ΔN_1-33_ mutant dissociated into dimeric molecules (Figs. 2G−I, Supplementary Fig. S4 and Table S5). In contrast, full-length *Rv*PrxL treated with 100 mM H_2_O_2_ undergoes only minor loop rearrangements, maintaining the overall 20-mer stacking assembly (Figs. 2J−K, Supplementary Figs. S3, S11 and Table S4). This local structural rearrangement in full-length *Rv*PrxL resembles the helical disruption of *Sm*Prx1 forming 20-mer assembly under an low-pH condition or when treated with highly concentrated H_2_O_2_ (31). These findings are consistent with Clear Native PAGE results (Supplementary Figs. S10C−D), which showed decomposition of high-oligomeric states induced by H_2_O_2_. These results suggest the critical role of the N-terminal region in stabilizing the high-oligomeric structure of *Rv*PrxL under oxidative conditions.

## Nuclear localization of *Rv*PrxL

Our cryo-EM analysis suggested that the N-terminal region of *Rv*PrxL may contribute to some physiological roles. To gain insight into the cellular function of *Rv*PrxL and its relationship with the N-terminal region, we next checked the cellular localization of *Rv*PrxL. A GFP-fused *Rv*PrxL (*Rv*PrxL–GFP) gene was introduced to HEK293T cells and we observed that *Rv*PrxL–GFP localizes exclusively to nucleus (Fig. 3A). In contrast, the ΔN_1–33_ mutant (ΔN_1–33_–GFP) localizes to cytoplasm, showing that the N-terminal region is necessary for nuclear localization.

**Fig. 3.**
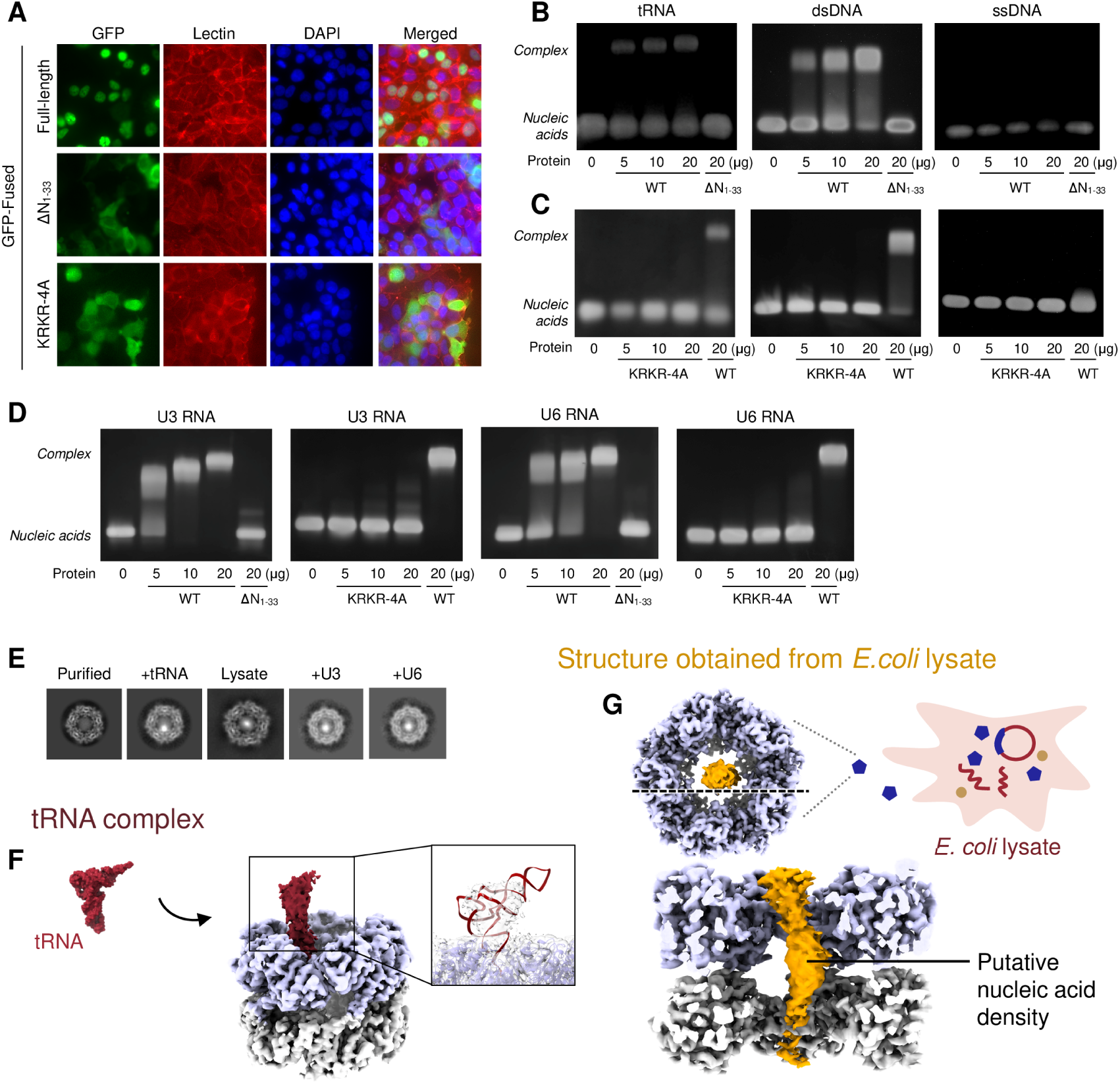
Nuclear localization and nucleic acid binding properties of *Rv*PrxL. (**A**) Expression of *Rv*PrxL–GFP, ΔN_1**–**33_–GFP and KRKR-4A–GFP in HEK293T cells. After the transfection, each cell was stained together with Lectin and DAPI. The experiment was repeated 2 times with similar results. (**B**) Nucleic acid binding of full-length and ΔN_1**–**33_ mutant of *Rv*PrxL. (**C**) Nucleic acid binding of KRKR-4A mutant of *Rv*PrxL. (**D**) U3 and U6 RNA binding of full-length and KRKR-4A mutant of *Rv*PrxL. (**E**) Representative 2D class images of *Rv*PrxL, tRNA complex, structure from *E. coli* lysate, U3 RNA complex and U6 RNA complex from cryo-EM analyses. (**F**) Cryo-EM structure of the tRNA complex. Left: Surface representation of the crystal structure of yeast phenylalanine tRNA (PDB: 1EHZ). Right: reconstructed cryo-EM map of the *Rv*PrxL–tRNA complex (*Rv*PrxL: purple and gray, tRNA: red). The inset shows superposition of the cryo-EM map (transparent surface) on tRNA (red cartoon) and *Rv*PrxL (purple cartoon) models. (**G**) Schematic illustration of the *E. coli* lysate sample and the cryo-EM map of the 20-mer observed in the lysate. The lower panel shows a sectional view along the dashed line in the upper-left panel. Putative nucleic acid-like density is colored in orange.

NLS Mapper (38) and NLSExplorer (39), which are web servers to predict nuclear localization signals (NLSs), show that *Rv*PrxL has an NLS in its N-terminal region (Supplementary Table S2), consistent with our results of the cellular localization experiment. It is noteworthy that the N-terminal region of *Rv*PrxL contains a KRKR motif (Fig. 1A), which is a well-known monopartite nuclear localization signal motif for many nuclear proteins including nuclear factor-kappa B (38, 56, 57).

To assess the role of this KRKR motif, we generated a mutant in which the KRKR motif is replaced with 4 alanine residues (KRKR-4A) and performed the same localization analysis. We observed that the KRKR-4A mutant localizes to cytoplasm (Fig. 3A), showing that the KRKR motif is crucial for transferring to the nucleus.

The KRKR motif is also found in the N-terminal region of several Prxs from anhydrobiotic tardigrades, *Hypsibius exemplaris* and *Paramacrobiotus metropolitanus*. In contrast, high-scoring hit sequences from other organisms identified by BLAST searches lack this motif (Supplementary Figs. S12A−B). Therefore, the KRKR motif may have been specifically acquired in the common ancestor of these tardigrades and retained as a lineage-specific feature. One of the tardigrade Prxs (OQV14346.1 from *H*. *exemplaris*) even possesses a zinc finger domain at its C-terminal region in addition to an N-terminal KRKR motif (Supplementary Fig. S12C), suggesting that certain tardigrade Prxs may have evolved to function in the nucleus.

## Nucleic acid binding properties of *Rv*PrxL

Given the positively charged nature of the tardigrade-specific N-terminal region and nuclear localization feature of *Rv*PrxL, we hypothesized that *Rv*PrxL interacts with nucleic acids. Electrophoretic mobility shift assays (EMSAs) clearly revealed that *Rv*PrxL binds to tRNA and dsDNA (Fig. 3B). Although a discrete shifted band was not apparent for ssDNA, the intensity of the unshifted ssDNA band decreased in a protein concentration–dependent manner, consistent with ssDNA binding. The absence of a visible shifted ssDNA band may reflect reduced fluorescence upon complex formation. For example, bound short ssDNA (47 nt) may be easily masked by the protein, limiting access of the fluorescent dye, and/or ssDNA captured within the protein assembly may adopt heterogeneous or flexible conformations that disfavor dye intercalation, thereby diminishing signal from the protein–ssDNA complex. These observations of EMSAs are consistent with the fact that careful purification processes were needed to completely eliminate contamination of nucleic acids from the protein sample (Supplementary Fig. S13).

By contrast, the ΔN_1–33_ mutant failed to induce a mobility shift even with plasmids, for which the full-length protein produced a band shift at low protein concentrations, indicating that the N-terminal region is necessary for the interaction with nucleic acids (Supplementary Fig. S14A).

We then analyze which amino acids in the N-terminal region is responsible for the binding ability. Because the KRKR motif seemed to be critical due to the positively charging nature, we employed the KRKR-4A mutant to check its ability of binding to nucleic acids. The KRKR-4A mutant did not show any shifts, as well as ΔN_1–33_ mutant, revealing that KRKR motif binds to nucleic acids (Fig. 3C and Supplementary Fig. S14B). This result strongly suggests that the other Prxs from tardigrade which contains the KRKR motif (Supplementary Fig. S12B) may also possess the nucleic acid binding ability. Recent studies have reported that Prxs interact with nucleic acids and protect them from nuclease degradation (58, 59). To investigate whether *Rv*PrxL possesses a similar protective property against dsDNA degradation, we performed a mobility shift assay under DNase exposure. In the control group without *Rv*PrxL, the intensity of the band significantly decreased upon DNase treatment (Supplementary Fig. S14C). In contrast, the bands were preserved in the *Rv*PrxL-treated group, albeit with reduced intensity after DNase exposure. The band intensity of remaining dsDNA exhibited an upward trend in the presence of *Rv*PrxL compared with the control (Supplementary Fig. S14D−E), demonstrating that *Rv*PrxL partially protects dsDNA from DNase-mediated degradation.

Because a previous study reports that in addition to the canonical enzymatic function, *Hs*Prx1 has the ability to bind small nucleolar RNAs (snoRNA) to function as an RNA chaperone and participate in the regulation of snoRNA (60), we tested whether *Rv*PrxL also interacts with tardigrade RNAs present in the nucleus. For this purpose, we used U3 snoRNA and U6 small nuclear RNA (snRNA) from *P. metropolitanus*, the reliably annotated RNA sequence of which can be obtained from the database. *P. metropolitanus* also has a Prx containing the tardigrade-specific N-terminal region with the KRKR motif (Fig. 1A, Supplementary Fig. S12B). Human U3 snoRNA is known to interact with *Hs*Prx1 (60) and is involved in the maturation of ribosomal RNAs. U6 snRNA is the most conserved RNA among eukaryotes (61) and is involved in splicing of mRNA precursors.

EMSAs using U3 and U6 RNAs show that wild-type *Rv*PrxL efficiently binds them while the ΔN_1–33_ and KRKR-4A mutants without the KRKR motif do not (Fig. 3D). Moreover, complete band shifts at 10–20 μg protein showed that *Rv*PrxL has higher affinity to these RNAs compared to tRNA. In the 20 μg protein conditions, the ΔN_1–33_ and KRKR-4A mutants showed a slight band shift or smearing of a band, implying non-specific binding or interaction through positively charged residues in the N-terminal region other than the KRKR motif. These findings suggest that *Rv*PrxL may also play a role in the regulation of RNAs in the tardigrade nucleus through the interaction at the N-terminal region, especially at the KRKR motif.

## Cryo-EM analyses visualized “on-ring” interaction with nucleic acids

Next, we sought to directly visualize the interaction between *Rv*PrxL and nucleic acids by cryo-EM, because, to our knowledge, no structural studies of Prx scaffolds in complex with nucleic acids have been reported to date. Initially, yeast tRNA was employed as a model nucleic acid, owing to its characteristic structural rigidity. At a 2D classification step, we found the existence of strong anomalous densities in the *Rv*PrxL ring, which was not observed in the 2D images of purified *Rv*PrxL (Fig. 3E). As a result, we could observe density attached to the N-terminal region of *Rv*PrxL after the 3D map reconstruction (3.40 Å resolution) (Fig. 3F, Supplementary Figs. S5, S15B, Table S5 and Video S1). Although we attempted local refinement focused on the tRNA-bound region, this did not substantially improve the map, likely because the tRNA density is small and conformationally heterogeneous. We therefore could not obtain a sufficiently clear map for atomic modeling of the tRNA region. Nevertheless, superposition of a tRNA model onto the cryo-EM map showed that the additional density overlaps with the model, supporting its interpretation as tRNA-associated density. Because the tRNA molecule sits on the 20-mer ring of *Rv*PrxL, we named this interaction manner “on-ring mode”. The diffuse density corresponding to tRNA suggests that the tRNA molecule is dynamic on *Rv*PrxL, likely reflecting transient and/or weak trapping.

## Cryo-EM visualization of “in-ring mode” complex by LApDoG approach

To examine *Rv*PrxL–nucleic acid complexes under more near-physiological conditions, we therefore tested an approach in which cell **<u>L</u>**ysates are **<u>Ap</u>**plied **<u>D</u>**irectly **<u>o</u>**n cryo-EM **<u>G</u>**rids (LApDoG), avoiding purification steps (Supplementary Fig. S2D). Although purification-free cryo-EM approaches, such as cryoPRISM (62) and *in-extracto* cryo-EM (63), have been reported, their targets have been limited to massive macromolecular complexes with high signal-to-noise ratios, such as ribosomes. In this study, we applied this approach to a smaller protein molecule. Because prokaryotic cells are simpler than eukaryotic cells and hence better suited for such direct, minimally processed observations of lysates, we used *E*. *coli* cells overexpressing *Rv*PrxL for cryo-EM analysis with the LApDoG approach. Recombinant eukaryotic proteins expressed in *E*. *coli* can form protein–DNA complexes in the bacterial cells, and the resulting complexes can closely resemble native structures (64).

The *E*. *coli* lysate containing many *Rv*PrxL molecules (Supplementary Fig. S2C) enabled successful collection of cryo-EM data by the LApDoG method. At a 2D classification step, we could observe the existence of anomalous densities in the *Rv*PrxL ring clearly, as observed in the tRNA complex data (Fig. 3E). While tRNA was bound off-center to the internal cavity, the density in the ring of this structure was located centrally in the cavity. Among 10-mer, 20-mer, 30-mer and 40-mer structures obtained, the 10-mer and 20-mer maps indicate the presence of nucleic acids in the central cavity (Supplementary Figs. S6, S15C−D and Table S6). Especially, the 20-mer map (3.21 Å resolution) most clearly shows putative nucleic acid density (Fig. 3G, Supplementary Video S2). This curved density was compatible to nucleic acid models even dsDNA (Supplementary Fig. S16). However, because this approach utilizes crude cell lysates without chromatographic purification, we were unable to determine the specific nucleic acid species corresponding to the observed putative density. Moreover, the ambiguity of the density hindered atomic model building, reflecting sequence non-specificity and/or conformational heterogeneity of the bound nucleic acids. Consistent with this idea, mobility shift assays also demonstrated the broad binding capacity of *Rv*PrxL to diverse nucleic acids, suggesting that the diffuse density reflects its intrinsic non-specific binding property.

The putative nucleic acid density of the lysate sample is in the 20-mer ring of *Rv*PrxL, so we named this interaction manner “in-ring mode”. This observation of the in-ring mode provides a structural context for our EMSA results with ssDNA, in which a discrete shifted band was not apparent despite evidence of binding. If single-stranded nucleic acids are captured within the ring, complex formation could reduce dye accessibility through protein shielding. In addition, conformational heterogeneity of the enclosed nucleic acids may disfavor dye intercalation, resulting in a weak or undetectable signal from the bound species.

## Cryo-EM analysis of U3 and U6 RNA complexes

We also performed cryo-EM analyses using U3 and U6 RNAs, which are expected to be more physiologically relevant partners. However, adding these RNAs to the *Rv*PrxL sample altered particle orientation distribution that limited the acquisition of sufficient side-view image of the particles. Although we could observe many particles showing putative RNA densities on raw micrographs and 2D classification images showing strong anomalous densities in the *Rv*PrxL ring (Fig. 3E and Supplementary Figs. S7−8), we could not obtain a 3D map that clearly shows RNA densities.

Formation of the in-ring complex may be favored if the oligomer transiently dissociates or exists as lower-order species before the reassembly around nucleic acids. Such remodeling of the oligomeric state could occur during importin-mediated nuclear transport, where passage through the nuclear pore may favor more compact or lower-order transport intermediates. In contrast, in our structural analyses, the *Rv*PrxL–RNA complexes were prepared *in vitro* by mixing RNAs with the preformed oligomeric protein. Under these conditions, *Rv*PrxL–RNA complexes would be expected to predominantly adopt the on-ring state, as observed in the tRNA-complex analysis. However, the conformational heterogeneity of U3 and U6 RNAs made it much more difficult to reconstruct a 3D map of the on-ring mode than in the case of the more rigid *Rv*PrxL–tRNA complex. We therefore performed deep 3D classification to search for less populated binding states. This analysis identified a small fraction of particles corresponding to an in-ring state, although the resulting map was of limited quality (Supplementary Figs. S7−8 and Table S5).

## Discussion

Our present study reveals that *Rv*PrxL represents an example of functional divergence within a Prx-like protein family in anhydrobiotic tardigrades. Although *Rv*PrxL retains an overall Prx architecture with high similarity to typical 2-Cys Prxs, it carries a glutamate residue at the position corresponding to Cys^P^. This substitution abolishes canonical peroxidase catalysis and chaperone activity. Additionally, our findings clearly showed that the tardigrade-specific KRKR motif drives the neo-functionalization of *Rv*PrxL, conferring both nuclear localization and nucleic acid-binding functions.

Our cryo-EM experiments revealed two distinct modes of nucleic-acid interaction with the Prx architecture, termed “on-ring” and “in-ring.” The coexistence of these modes suggests that *Rv*PrxL can engage nucleic acids through more than one binding pathway. One possibility is that the two modes reflect differences in the order of oligomer assembly and nucleic-acid binding (Fig. 4, upper right panel). In the on-ring mode, preassembled *Rv*PrxL oligomers may capture nucleic acids on the outer surface of the ring, whereas in the in-ring mode, lower-order *Rv*PrxL species may reassemble around nucleic acids, resulting in their incorporation into the central cavity. These alternative pathways could secondarily influence which nucleic-acid structures or conformational states are accommodated, thereby allowing *Rv*PrxL to support multiple modes of nucleic-acid protection or regulation.

**Fig. 4.**
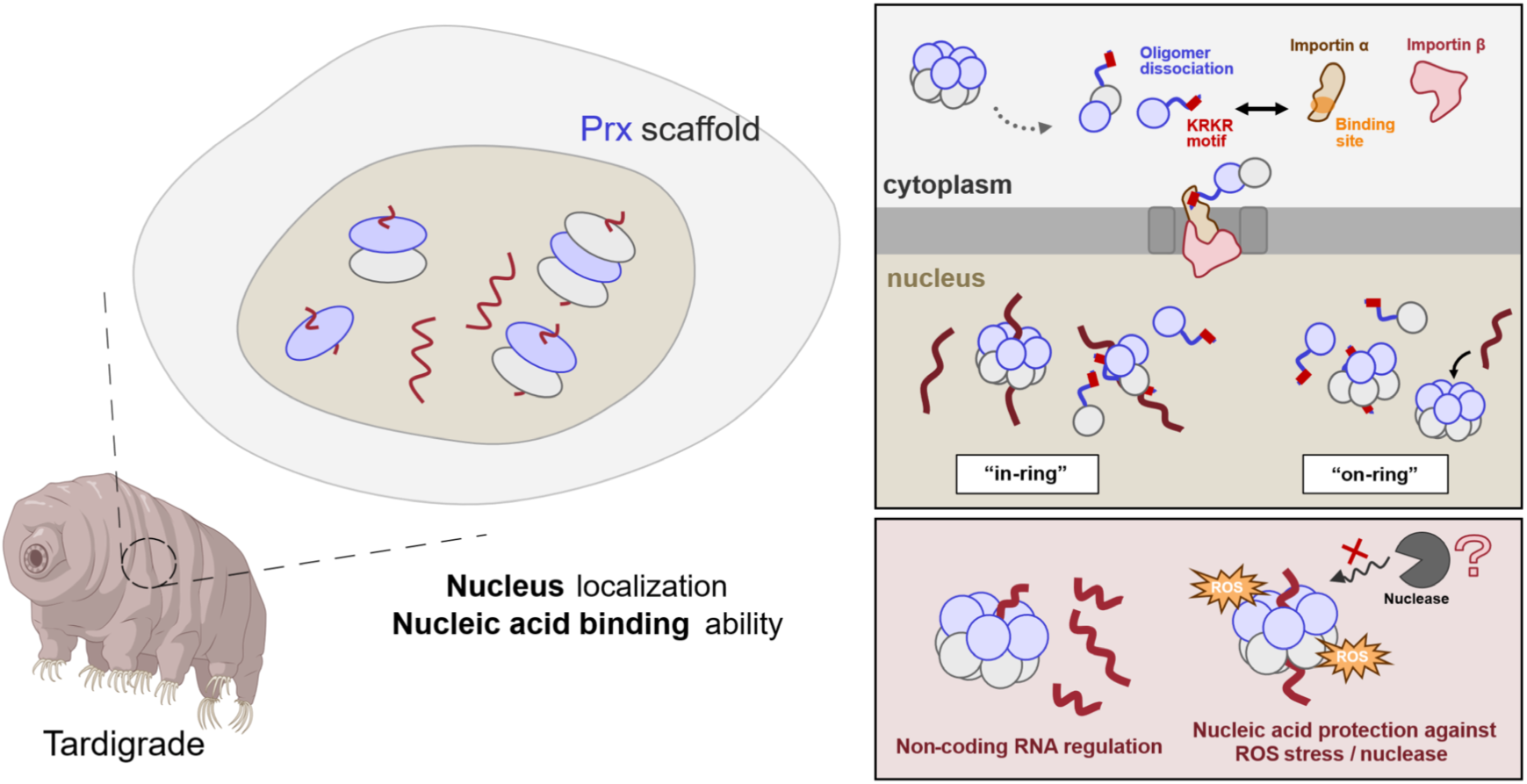
Possible physiological functions and a proposed mechanistic model of neofunctionalized tardigrade Prxs. *Rv*PrxL may form the “in-ring” mode when it reassembles around nucleic acids in the nucleus, whereas preassembled 20-mers may bind nucleic acids in the “on-ring” mode (upper right panel). These two modes may therefore reflect co-assembly with nucleic acids and post-assembly binding, respectively. Tardigrade Prxs that have evolved to localize to the nucleus may protect nucleic acids against ROS stresses or nuclease, or may contribute to the regulation of non-coding RNAs (lower right panel). Some figures are from BioRender (https://www.biorender.com).

The assembly-dependent nucleic-acid binding model is further supported by the difference between the *in vitro* reconstitution and LApDoG conditions. In the *in vitro* reconstitution experiments, RNAs were mixed with preformed *Rv*PrxL oligomers, a condition that would favor post-assembly binding and is consistent with the predominant observation of the on-ring mode. By contrast, in the LApDoG analysis, nucleic-acid density was observed mainly within the central cavity of the *Rv*PrxL oligomer. Because *Rv*PrxL and endogenous nucleic acids could have encountered each other during protein folding and/or oligomerization, this condition may allow nucleic acids to be incorporated during or before completion of oligomer assembly. Thus, the predominance of the in-ring mode in LApDoG correlates well with a co-assembly pathway, whereas the on-ring mode observed after *in vitro* mixing is more consistent with post-assembly binding. Moreover, because putative nucleic acid density was retained after overnight dialysis in LApDoG, the enclosed nucleic acid appears to be protected, at least in part, from endogenous nucleases present in the lysate.

We speculate that the in-ring mode represents a sequestration state in which nucleic acids are enclosed within the oligomeric assembly, thereby limiting their accessibility to other macromolecules such as nucleases and possibly buffering them from damage under stress. The in-ring structure may therefore offer a structural explanation for previous reports that some Prxs protect nucleic acids from nuclease degradation (58, 59). Consistent with this idea, our DNase protection assay showed that *Rv*PrxL increased the amount of dsDNA remaining after DNase treatment, although the protection was incomplete. This incomplete protection is compatible with the coexistence of multiple binding modes including the on-ring mode, which may be more readily formed when nucleic acids bind to preassembled *Rv*PrxL oligomers. In contrast to the enclosed in-ring state, the weaker on-ring mode may correspond to a transient capture state that enables rapid sampling, handoff, or localized recruitment of nucleic acids without full enclosure. In this view, in-ring binding would be suited for protection or storage, whereas on-ring binding could facilitate dynamic regulatory interactions (Fig. 4, lower right panel).

Due to the limitations of our study, we cannot discuss details of the physiological function of this protein. However, a transcriptome analysis shows that *Rv*PrxL is most abundant in the embryonic stage (12), implying that *Rv*PrxL may protect nucleic acids from damage that occurs during cell division and growth, or may contribute to the regulation of non-coding RNAs at the developmental stage. Specifically, the fact that the tardigrade-specific N-terminal region helps to retain the quaternary structure even in the presence of high concentration of H_2_O_2_ suggests that *Rv*PrxL can function under severe oxidative stress conditions and its rigid structure enables protection of captured nucleic acids from oxidative stress.

Nucleic-acid binding by Prx-family proteins has also been reported in humans and other organisms (58, 59, 60, 65), yet the structural basis of these interactions, which is needed for the deeper understanding of this phenomenon, remains poorly defined. Our work therefore provides a framework for thinking about how a Prx scaffold can engage nucleic acids and may motivate comparable analyses in other systems. In particular, further application of the LApDoG approach to capture structures under intracellular-like conditions could help clarify physiologically relevant nucleic-acid interaction modes. It has been widely assumed that anhydrobiotic tardigrades partially gain extreme stress tolerance by expanding antioxidant enzymes through gene duplication (12, 18). Our findings argue that this prevailing view is incomplete because expansion of antioxidant gene families may also conceal extensive functional diversification, including paralogs that no longer operate as canonical detoxifying enzymes. Consistent with this broader interpretation, one of superoxide dismutases from *R*. *varieornatus* has recently been reported to be non-catalytic (66), supporting the idea that loss of enzymatic activity may enable the emergence of alternative roles within expanded antioxidant families. A plausible reason why such repurposing is particularly feasible in anhydrobiotic tardigrades is their redundant H₂O₂-defense system. For example, beyond Prx expansion, some tardigrades show an expansion of catalases *via* horizontal gene transfer (12, 18, 67). This multilayered redundancy (68–70) could buffer fitness costs when an individual Prx paralog loses canonical peroxidase function, allowing the Prx fold to be explored as a scaffold for neofunctionalization rather than being maintained solely as an enzymatic backup.

We propose a stepwise scenario for the evolution of *Rv*PrxL. The N-terminal extension may first have been retained because it increased structural robustness, helping the ring-shape assembly remain intact under oxidative stress conditions. Subsequently, or potentially in parallel, the KRKR motif may have evolved to confer nuclear targeting, opening opportunities for interactions with molecules in the nucleus. Once a non-enzymatic role became established and the ability of H_2_O_2_ detoxification was buffered by redundancy of other antioxidant enzymes, selective pressure to maintain canonical Prx catalysis could have weakened, permitting decay of the catalytic function.

## Acknowledgements

We thank staff at the Scientific Imaging Section of Okinawa Institute of Science and Technology for cryo-EM training and preliminary observation. We thank Mr. Hiroki Tanino and Nao Arakawa and Ms. Yuki Shimokawa (Graduate School of Pharmaceutical Sciences, the University of Osaka) for their support in cryo-EM data collection. The authors also thank the lab members of Tsuyoshi Inoue lab for insightful discussions and continuous support throughout the study. The authors used ChatGPT and Google Gemini to assist with English language editing and grammatical improvement. All AI-assisted text was reviewed and edited by the authors, who take full responsibility for the final content of the manuscript.

## Author Contributions

Haruka Yamato: Conceptualization, Investigation, Formal analysis, Methodology, Validation, Visualization, Writing—original draft, Writing—review & editing. Yohta Fukuda: Conceptualization, Investigation, Formal analysis, Methodology, Validation, Visualization, Supervision, Funding acquisition, Writing—original draft, Writing—review & editing. Masanori Obana: Investigation, Visualization, Writing—review & editing. Yasushi Fujio: Investigation, Writing—review & editing. Tsuyoshi Inoue: Investigation, Funding acquisition, Writing—review & editing.

## Data Availability Statement

The data that support the findings of this study are available on request from the corresponding author. The cryo-EM density maps and the atomic coordinates of full-length *Rv*PrxL, full-length *Rv*PrxL treated with 100 mM H_2_O_2_, ΔN_1–33_ mutant, tRNA−*Rv*PrxL complex, *Rv*PrxL obtained from unpurified *E. coli* lysate (10-mer, 20-mer, 30-mer and 40-mer) have been deposited in the Electron Microscopy Data Bank (EMDB) and the PDB with EMDB accession codes EMD-69157, EMD-69159, EMD-69248, EMD-69249, EMD-69250, EMD-69251, EMD-69252 and EMD-69253 and PDB accession codes 23PU, 23PV and 23TW, respectively. The raw cryo-EM datasets used to determine these deposited structures have been deposited in EMPIAR (https://www.ebi.ac.uk/empiar/). Fluorescence microscopy images obtained from the subcellular localization experiments have been deposited in the Systems Science of Biological Dynamics database (SSBD) repository under accession code ssbd-repos-000508.

