## Supplementary Data for "Cryo-EM structures of a neofunctionalized tardigrade peroxiredoxin specialized for nucleic acid binding"

**Supplementary Tables**

Table S1 The list of peroxiredoxins and GenBank accessions.

Table S2 NLS prediction with NLS Mapper and NLSExplorer.

Table S3 Primer sequences used to amplitude *Rv*PrxL.

Table S4 Cryo-EM data collection, refinement, and validation statistics.

Table S5 Cryo-EM data collection statistics of unmodeled structures.

Table S6 Cryo-EM data collection statistics of the structure obtained from *E. coli* lysate.

Table S7 DNA and RNA sequences used in mobility shift assay.

**Supplementary Figures**

Fig. S1 Sequence alignment of *Rv*PrxL with *Hs*Prx3 and *Sm*Prx1.

Fig. S2 Sample preparation and schematic of LApDoG method.

Fig. S3 Single-particle cryo-EM processing workflow and reconstructions of full-length *Rv*PrxL.

Fig. S4 Single-particle cryo-EM processing workflow and reconstructions of $\Delta N_{1-33}$ mutant.

Fig. S5 Single-particle cryo-EM processing workflow and reconstructions of tRNA complex.

Fig. S6 Single-particle cryo-EM processing workflow and reconstructions of structure obtained from *E. coli* lysate overexpressing *Rv*PrxL.

Fig. S7 Single-particle cryo-EM processing workflow and reconstructions of U3 RNA complex.

Fig. S8 Single-particle cryo-EM processing workflow and reconstructions of U6 RNA complex.

Fig. S9 *Rv*PrxL activity assays and structural analysis of full-length *Rv*PrxL.

Fig. S10 Structural analysis of $\Delta N_{1-33}$ mutant and clear native PAGE.

Fig. S11 Structural analysis of $H_2O_2$-treated *Rv*PrxL.

Fig. S12 Tardigrade-specific KRKR motif revealed from comparative analysis of Prx homologs.

Fig. S13 *Rv*PrxL purification process and UV-vis absorption spectroscopy.

Fig. S14 Mobility shift assay with plasmid and nuclease protection assay.

Fig. S15 Low-threshold maps and cryo-EM maps of *Rv*PrxL structure obtained from *E.coli* lysate overexpressing *Rv*PrxL.

Fig. S16 Superposition of DNA model to putative nucleic acid density.

**Supplementary Videos**

Video S1 Cryo-EM map of the tRNA complex.

Video S2 Cryo-EM map of a 20-mer with putative nucleic acid density.

**Supplementary Tables**

**Table S1 | The list of peroxiredoxins and GenBank accessions.**

| Taxon name | GenBank accession |
|---|---|
| ***Ramazzottius varieornatus* [*Rv*PrxL]** | **GAV04299.1** |
| *Homo Sapiens* [*Hs*Prx1] | NP_001189360.1 |
| *Homo Sapiens* [*Hs*Prx3] | NP_006784.1 |
| *Homo Sapiens* [*Hs*Prx4] | NP_006397.1 |
| *Schistosoma mansoni* [*Sm*Prx1] | 3ZVJ_A |
| *Paramacrobiotus metropolitanus* [*Pm*PrxL] | XP_055327524.1 |
| *Hypsibius exemplaris* [*He*Prx4] | OQV14348.1 |
| *Varroa destructor* [*Vd*PrxL] | XP_022651311.1 |
| *Adineta vaga* | UJR36090.1 |
| *Priapulus caudatus* | XP_014667732.1 |
| *Paralvinella palmiformis* | KAK2149584.1 |
| *Plutella xylostella* | XP_011561417.3 |
| *Capitella teleta* | ELU16156.1 |
| *Watersipora subatra* | XP_067951445.1 |
| *Tropilaelaps mercedesae* | OQR78401.1 |
| *Brachionus plicatilis* | RNA09967.1 |
| *Holothuria leucospilota* | KAJ8027160.1 |
| *Eurosta solidaginis* | XP_067643741.1 |
| *Zaprionus bogoriensis* | KAH8398855.1 |
| *Gigantopelta aegis* | XP_041352234.1 |
| *Adineta ricciae* | CAF1146296.1 |
| *Didymodactylos carnosus* | CAF1548135.1 |
| *Macrostomum lignano* | PAA87609.1 |
| *Amphiura filiformis* | XP_072035997.1 |
| *Lineus longissimus* | XP_064629193.1 |
| *Solemya velum* | KAL5010482.1 |
| *Antedon mediterranea* | XP_071954890.1 |
| *Hypsibius exemplaris*<br>(zinc finger domain containing Prx) | OQV14346.1 |

**Table S2 | NLS prediction with NLS Mapper and NLSExplorer.**

| **NLS Mapper** | |
|---|---|
| **Sequence** | **Score** |
| SSFKRKRAD | 8 |
| SSFKRKRADS | 6 |

| **NLSExplorer** | | |
|---|---|---|
| **Sequence** | **Recommendation score** | **Entropy score** |
| SFKRKRADSPQPG | 0.98579 | 3.08506 |

**Table S3 | Primer sequences used to amplitude *Rv*PrxL.**

| **Construct** | | **Primer Sequence** |
|---|---|---|
| Full-length | Forward | 5' ATGTCGAGCTTCAAGCGAAAGCGTGCGGATTCTCCA CAGC 3' |
| | Reverse | 5' TTATTTGTGGTGCATCTTGAAATACTCTCG 3' |
| $\Delta N_{1-33}$ mutant | Forward | 5' GCTGGTGATCATCGCATTTACACAT 3' |
| | Reverse | 5' CATGGTATATCTCCTTCTTAAAGTT 3' |

**Table S4 | Cryo-EM data collection, refinement, and validation statistics.**

| | **Full-length** | **Full-length + $H_2O_2$** | **$\Delta N_{1\text{–}33}$ mutant** |
|---|---|---|---|
| PDB ID | 23PU | 23PV | 23TW |
| EMDB ID | EMD-69157 | EMD-69159 | EMD-69248 |
| **Data collection and processing** | | | |
| Magnification | 60,000 | 60,000 | 60,000 |
| Voltage (kV) | 200 | 200 | 200 |
| Electron exposure (e-/Å$^2$) | 40.00 | 40.00 | 40.00 |
| Defocus range (μm) | −0.7 to −2.2 | −0.7 to −2.2 | −0.7 to −2.2 |
| Pixel size (Å) | 0.83 | 0.83 | 0.83 |
| Symmetry imposed | *D5* | *D5* | *C5* |
| Initial particle images (no.) | 3,276,108 | 2,050,898 | 2,470,287 |
| Final particle images (no.) | 177,997 | 279,910 | 557,497 |
| Map resolution (Å) | 2.65 | 2.96 | 3.02 |
| FSC threshold | 0.143 | 0.143 | 0.143 |
| **Refinement** | | | |
| Initial model used | Alphafold2 | Full-length | Full-length |
| Model resolution (Å) | 2.8 | 3.3 | 3.3 |
| FSC threshold | 0.5 | 0.5 | 0.5 |
| Map sharpening *B*-factor (Å$^2$) | 109.8 | 122.0 | 87.7 |
| Model composition | | | |
| Non-hydrogen atoms | 2745 | 2721 | 2693 |
| Protein residues | 351 | 348 | 344 |
| Water | 0 | 0 | 0 |
| Ligands | 0 | 0 | 0 |
| *B*-factors (Å$^2$) | | | |
| Protein | 41.71 | 55.21 | 143.53 |
| Ligand | - | - | - |
| R.m.s. devations | | | |
| Bond lengths (Å) | 0.004 (0) | 0.005 (0) | 0.003 (0) |
| Bond angles (°) | 0.827 (8) | 1.053 (7) | 0.591 (1) |
| Validation | | | |
| MolProbity score | 1.69 | 1.55 | 1.66 |
| Clashscore | 6.08 | 4.08 | 6.74 |
| Ramachandran plot (%) | | | |
| Favored | 96.25 | 94.77 | 95.88 |
| Allowed | 3.75 | 5.23 | 4.12 |
| Outliers | 0.00 | 0.00 | 0.00 |
| Model vs Map | | | |
| CC (mask) | 0.86 | 0.86 | 0.91 |
| CC (main chain) | 0.87 | 0.87 | 0.88 |
| CC (side chain) | 0.85 | 0.85 | 0.86 |

**Table S5 | Cryo-EM data collection statistics of unmodeled structures.**

| | **$\Delta N_{1-33}$ mutant + $H_2O_2$** | **tRNA complex** | **U3 RNA complex** | **U6 RNA complex** |
|---|---|---|---|---|
| EMDB ID | **-** | EMD-69249 | - | - |
| **Data collection and processing** | | | | |
| Magnification | 60,000 | 60,000 | 60,000 | 60,000 |
| Voltage (kV) | 200 | 200 | 200 | 300 |
| Electron exposure (e-/Å$^2$) | 40.00 | 40.00 | 40.00 | 40.00 |
| Defocus range (µm) | −0.7 to −2.2 | −0.7 to −2.2 | −0.7 to −2.2 | −0.6 to −1.6 |
| Pixel size (Å) | 0.83 | 0.83 | 0.83 | 0.86 |
| Symmetry imposed | *C1* | *C1* | *C1* | *C1* |
| Initial particle images (no.) | 3,117,531 | 7,641,544 | 3,043,627 | 15,790,961 |
| Final particle images (no.) | 101,782 | 109,368 | 5,768 | 13,328 |
| Map resolution (Å) | - | 3.40 | 5.98 | 4.46 |
| FSC threshold | 0.143 | 0.143 | 0.143 | 0.143 |

**Table S6 | Cryo-EM data collection statistics of the structure from *E. coli* lysate.**

| | **10-mer** | **20-mer** | **30-mer** | **40-mer** |
|---|---|---|---|---|
| EMDB ID | EMD-69250 | EMD-69251 | EMD-69252 | EMD-69253 |
| **Data collection and processing** | | | | |
| Magnification | 60,000 | 60,000 | 60,000 | 60,000 |
| Voltage (kV) | 200 | 200 | 200 | 200 |
| Electron exposure (e-/Å$^2$) | 40.00 | 40.00 | 40.00 | 40.00 |
| Defocus range (µm) | −0.7 to −2.2 | −0.7 to −2.2 | −0.7 to −2.2 | −0.7 to −2.2 |
| Pixel size (Å) | 0.83 | 0.83 | 0.83 | 0.83 |
| Symmetry imposed | *C1* | *C1* | *C1* | *C1* |
| Initial particle images (no.) | 5,821,219 | | | |
| Final particle images (no.) | 7,304 | 62,035 | 44,749 | 45,724 |
| Map resolution (Å) | 4.19 | 3.21 | 3.19 | 3.21 |
| FSC threshold | 0.143 | 0.143 | 0.143 | 0.143 |

**Table S7 | DNA and RNA sequences used in mobility shift assay.**

[1] A primer made and used in our previous study (Sassa and Yamato *et al. JBC* 2025)

| | | **DNA sequence** |
|---|---|---|
| **ssDNA**[1] | | 5’ GAAAACCTGTATTTTCAGGGCATGGAAACCAAAACCGAAACCAAAAC 3’ |
| **dsDNA** | | 5’ GAAAACCTGTATTTTCAGGGCATGGAAACCAAAACCGAAACCAAAAC 3’ |
| ***P. metropolitanus*** **small nucleolar U3 RNA** (XR_008690481.1) | | 5’ AGCGACCUACUUCACAGGAUCAGUGCAUUAGGUUGUUAACUGCGAACAGUCUGGAACUGUGCGCGUACACCAAACCUCGAUGAUGAGGAGUAGUGAUCCCUUCUGAGCGCGAAGCCGUUUUGGGCGGUUGGUCGAAAGACUGUCGCCCUGACAUUGAUGACCGUUCCUUUCCCCCUUGACUACGCGAUAGUCGGGGGAUGAGGAGGGAGGGAUCGCAUUCCGAGCGGU 3’ |
| ***P. metropolitanus*** **U6 spliceosomal RNA** (XR_008689789.1) | | 5’ GAGGUGUUUCCACCUCAUAUACUAAAUUGGAAACGAUACAGAGAAGAUUAGCAUGGCCCCUGCGCAAGGAUGACACGCAAAAUCGUGAAGCGUUUCCAAAUUUU 3’ |
| U3 RNA primer | Forward | 5’ CGCGAAATTAATACGACTCACTATAGGGAGCG 3’ |
| | Reverse | 5’ ACCGCTCGGAATGCGATCCC 3’ |
| U6 RNA primer | Forward | 5’ CGCGAAATTAATACGACTCACTATAGGGGAGG 3’ |
| | Reverse | 5’ AAAATTTGGAAACGCTTCACGATTTTGCGTG 3’ |

### Supplementary Figures

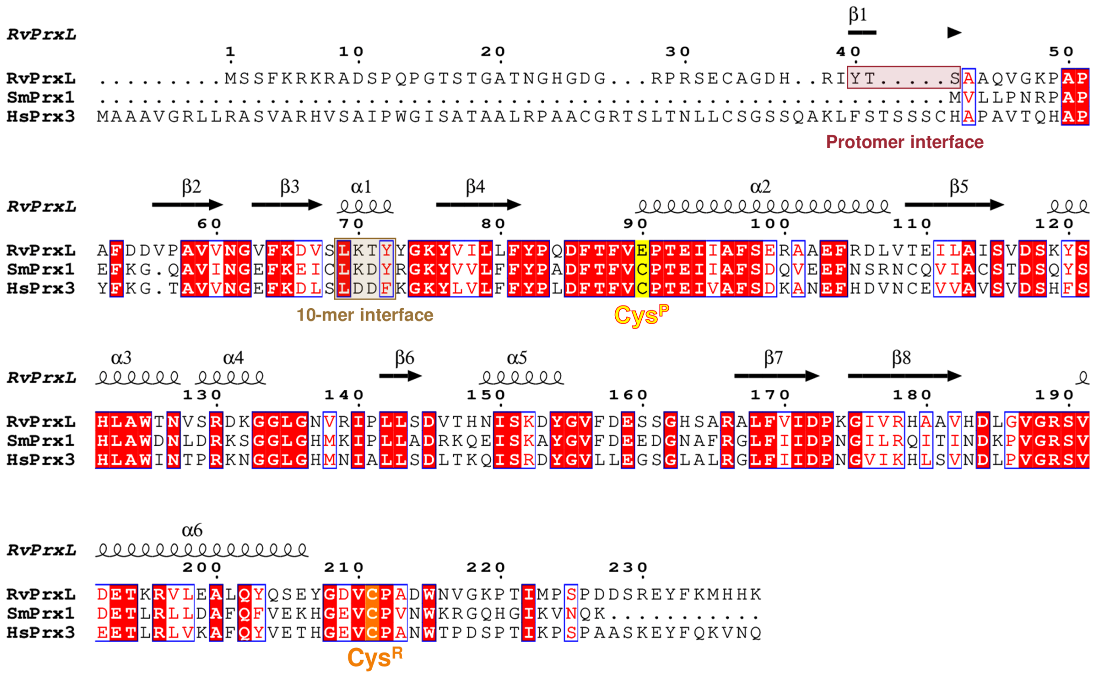

**Fig. S1 Sequence alignment of *Rv*PrxL with *Hs*Prx3 and *Sm*Prx1.**

The secondary structure of molA from the *Rv*PrxL is shown above the alignment. The $Cys^P$ (peroxidatic cysteine) and $Cys^R$ (resolving cysteine) are highlighted. The 10-mer interface region and protomer interface region are indicated by beige box and red box, respectively. Notably, the N-terminal region of the *Hs*Prx3 sequence (1–61) is known to be proteolytically cleaved *in vivo* as a mitochondrial signal peptide. *Rv*PrxL shows 56.3% sequence identity with *Hs*Prx3 and 54.9% sequence identity with *Sm*Prx1.

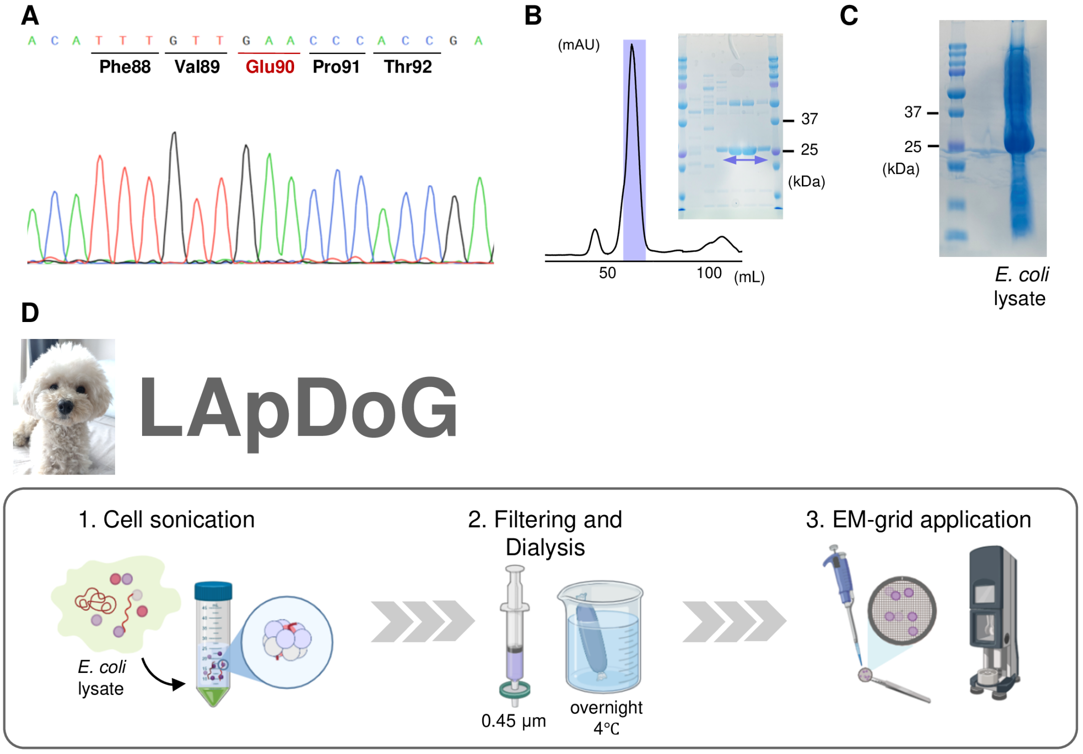

**Fig. S2 Sample preparation and schematic of LApDoG method.**

(**A**) A representative portion of the sequencing data visualized by SnapGene. (**B**) The result of size exclusion column chromatography and SDS-PAGE during the purification of Full-length *Rv*PrxL (monomer: 26 kDa). The area indicated in purple bar and arrow was collected as a final sample. (**C**) SDS-PAGE of *E. coli* lysate overexpressing *Rv*PrxL. (**D**) Schematic of the LApDoG method. Figures are from BioRender (https://www.biorender.com). The image of the lapdog, Jewel, is provided as a courtesy of Mr. Yasuhiro Ashida.

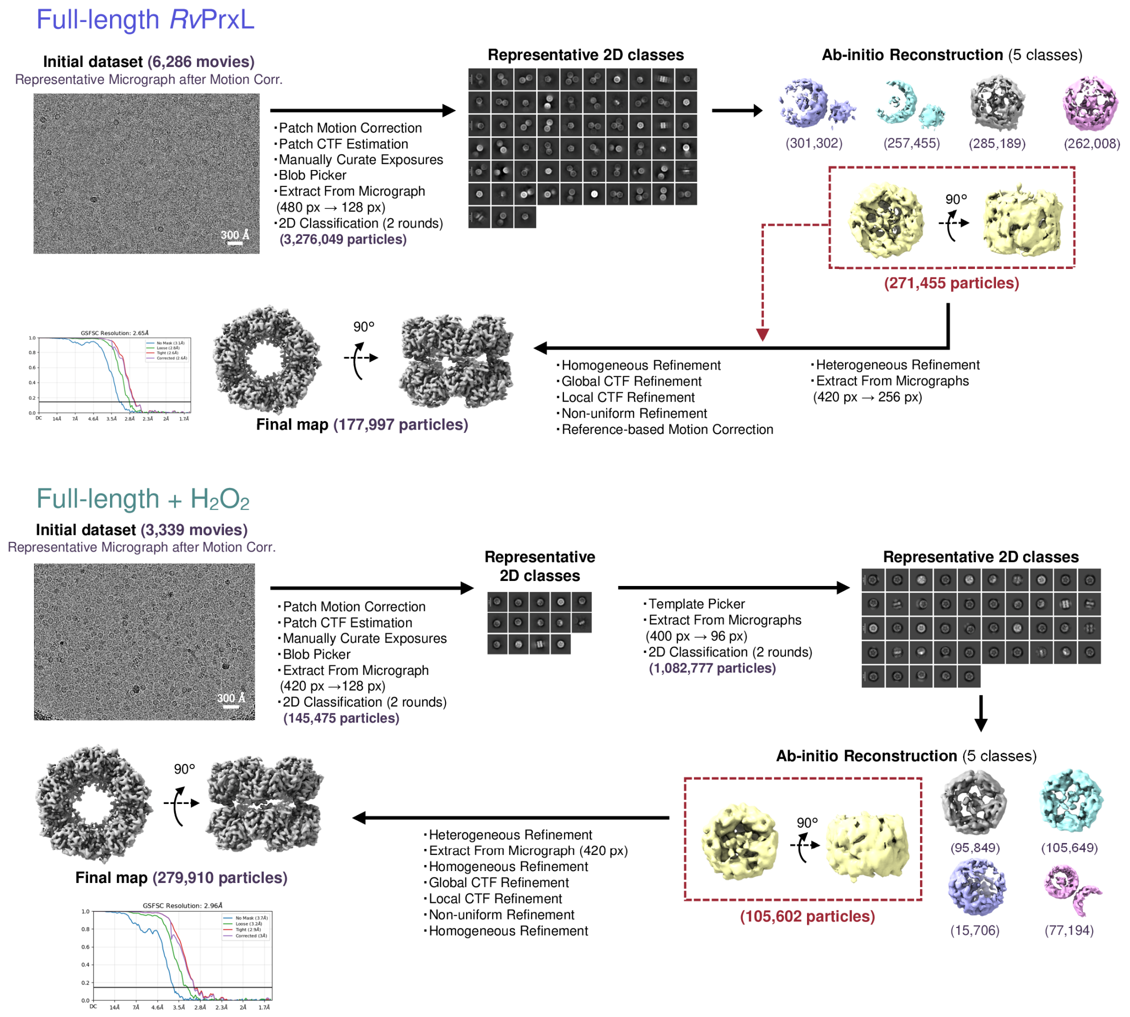

**Fig. S3 Single-particle cryo-EM processing workflow and reconstructions of full-length *Rv*PrxL.**

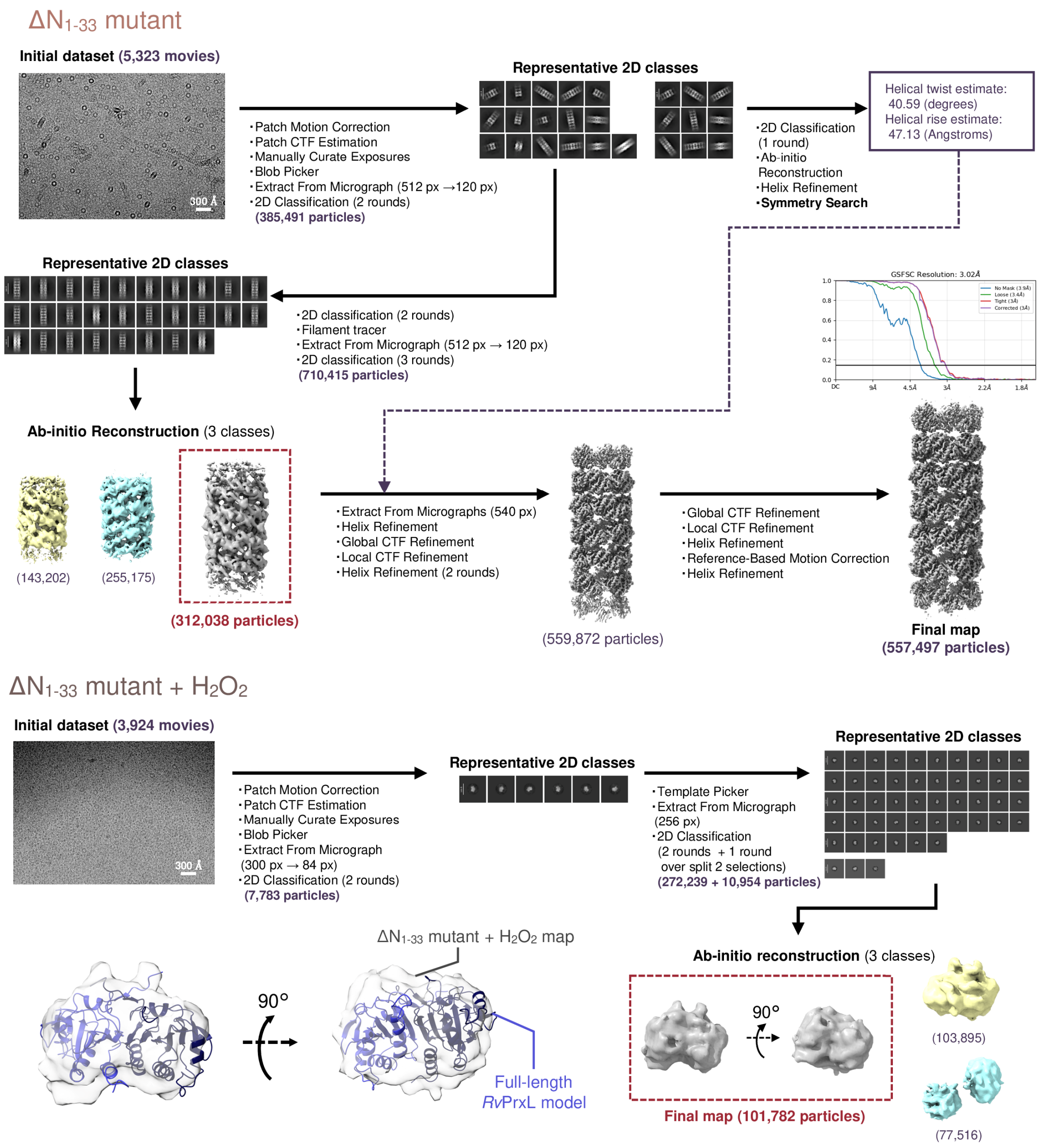

**Fig. S4 Single-particle cryo-EM processing workflow and reconstructions of $\Delta N_{1-33}$ mutant.**

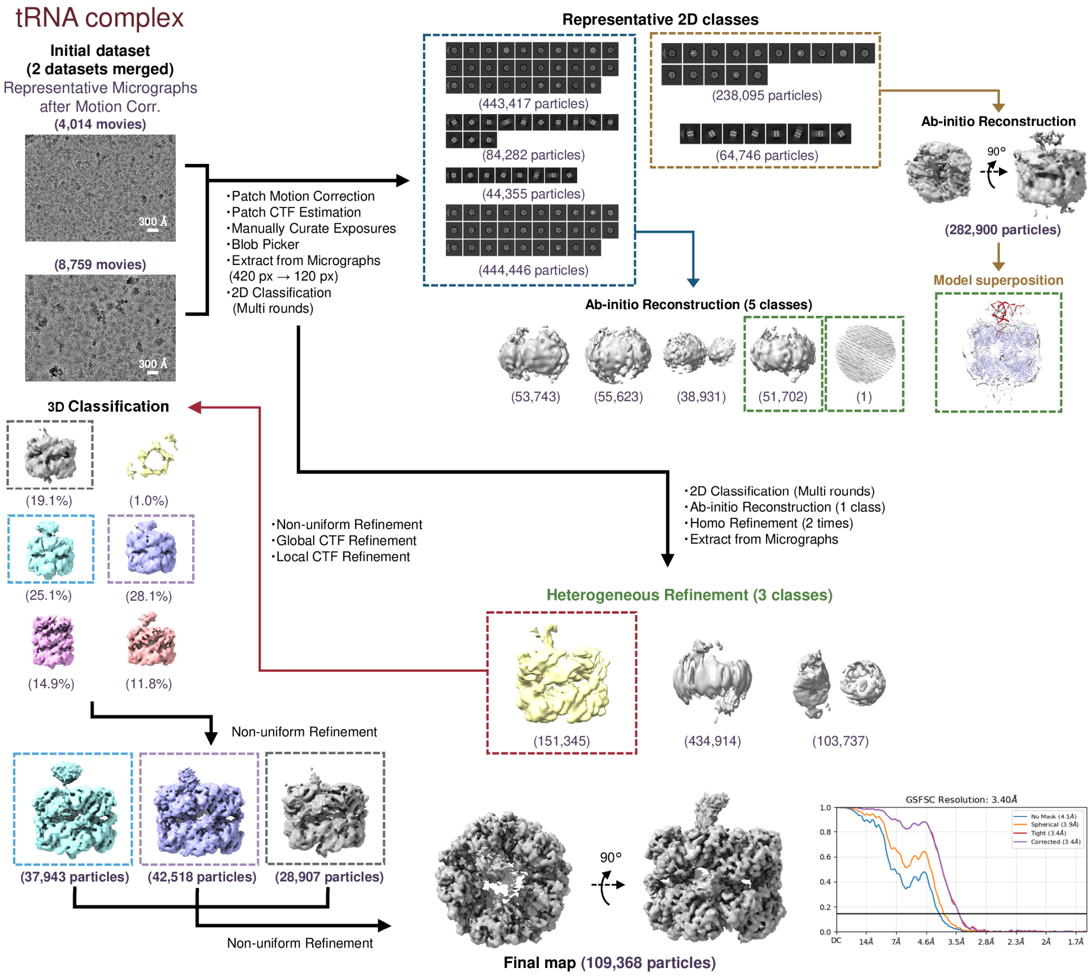

**Fig. S5 Single-particle cryo-EM processing workflow and reconstructions of tRNA complex.** The maps indicated in the dotted green lines were used as initial templates of Heterogeneous Refinement.

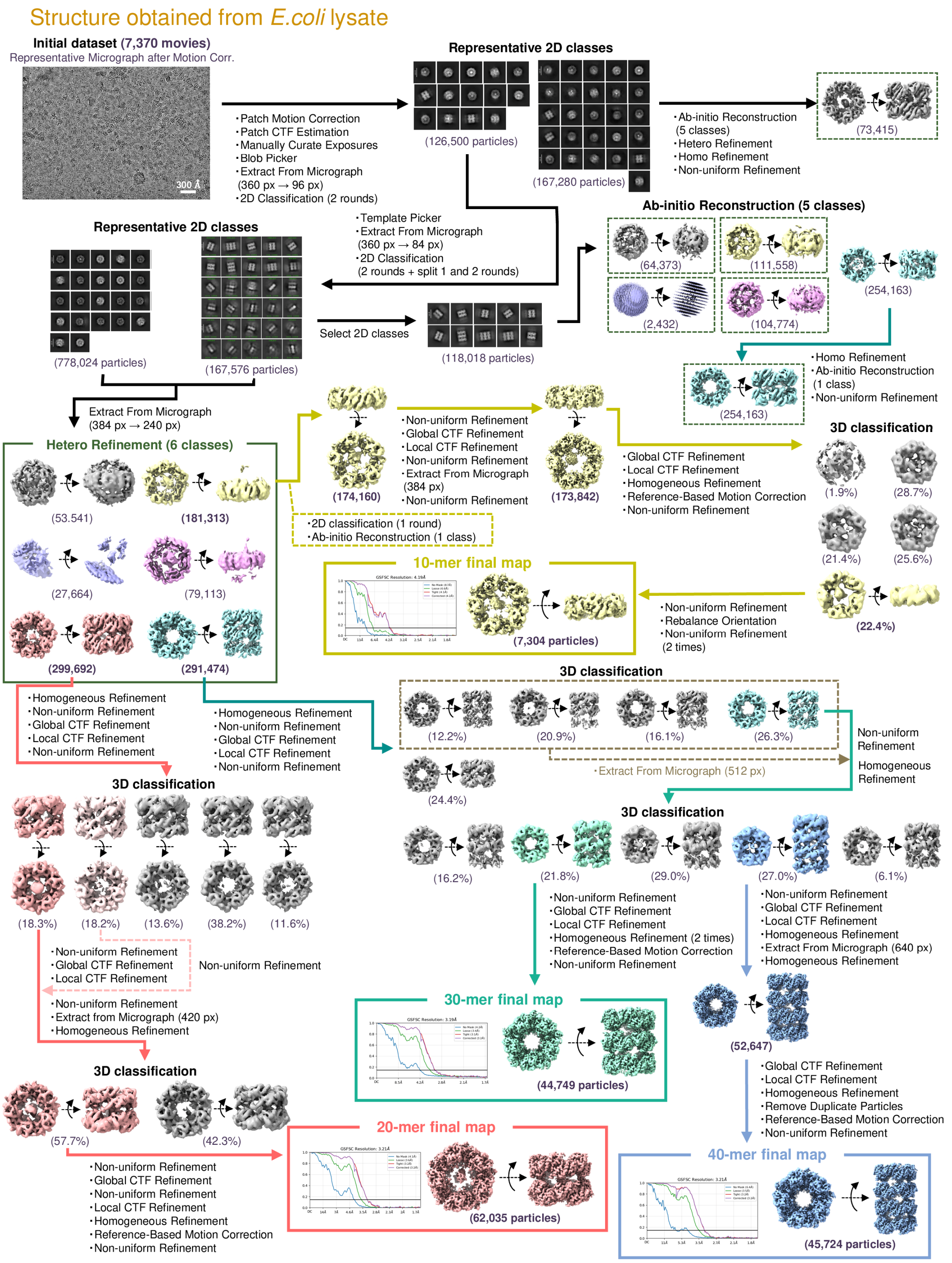

**Fig. S6 Single-particle cryo-EM processing workflow and reconstructions of structure obtained from *E. coli* lysate overexpressing *Rv*PrxL.**

The maps indicated in the dotted green lines were used as initial templates of Heterogeneous Refinement.

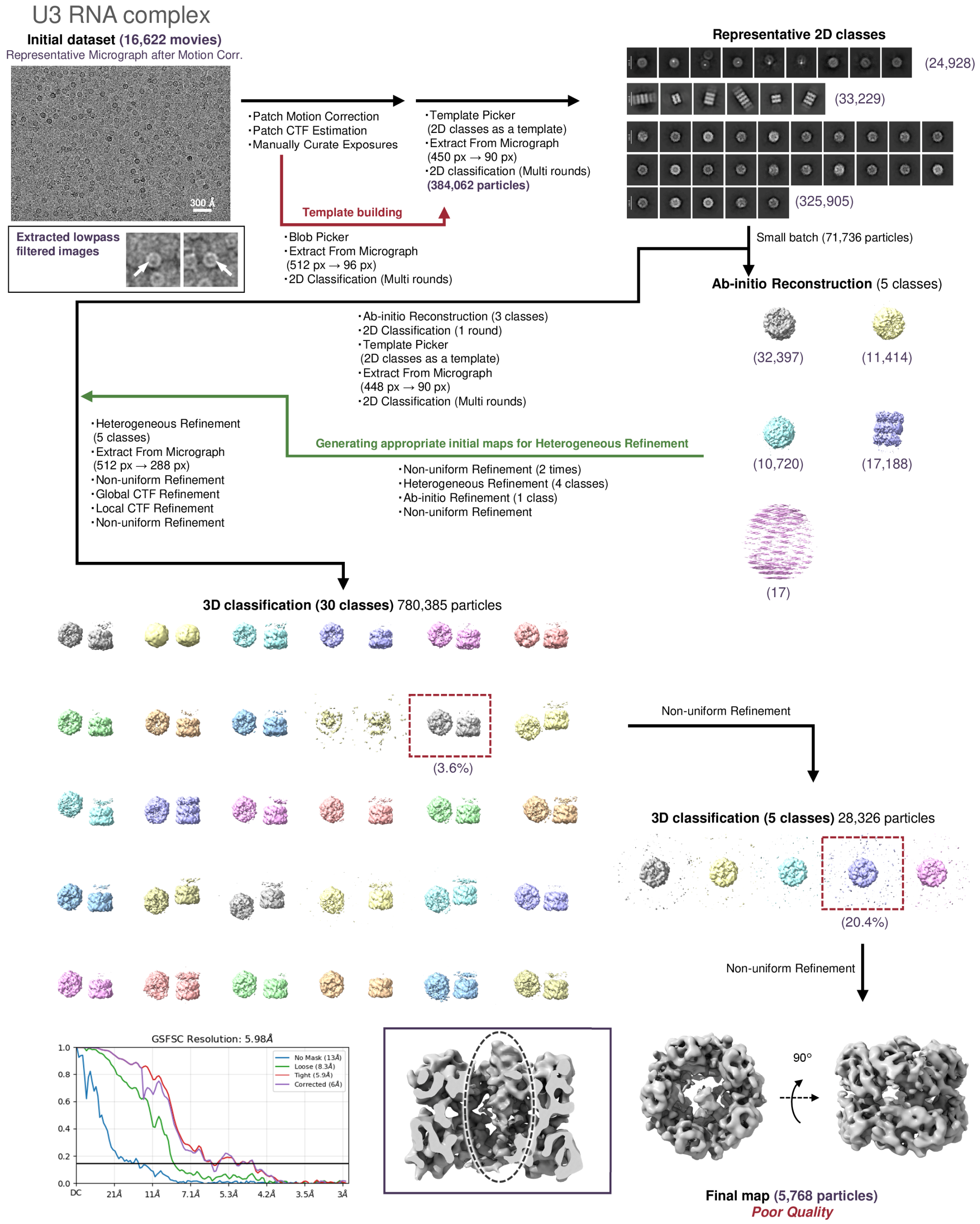

**Fig. S7 Single-particle cryo-EM processing workflow and reconstructions of U3 RNA complex.**

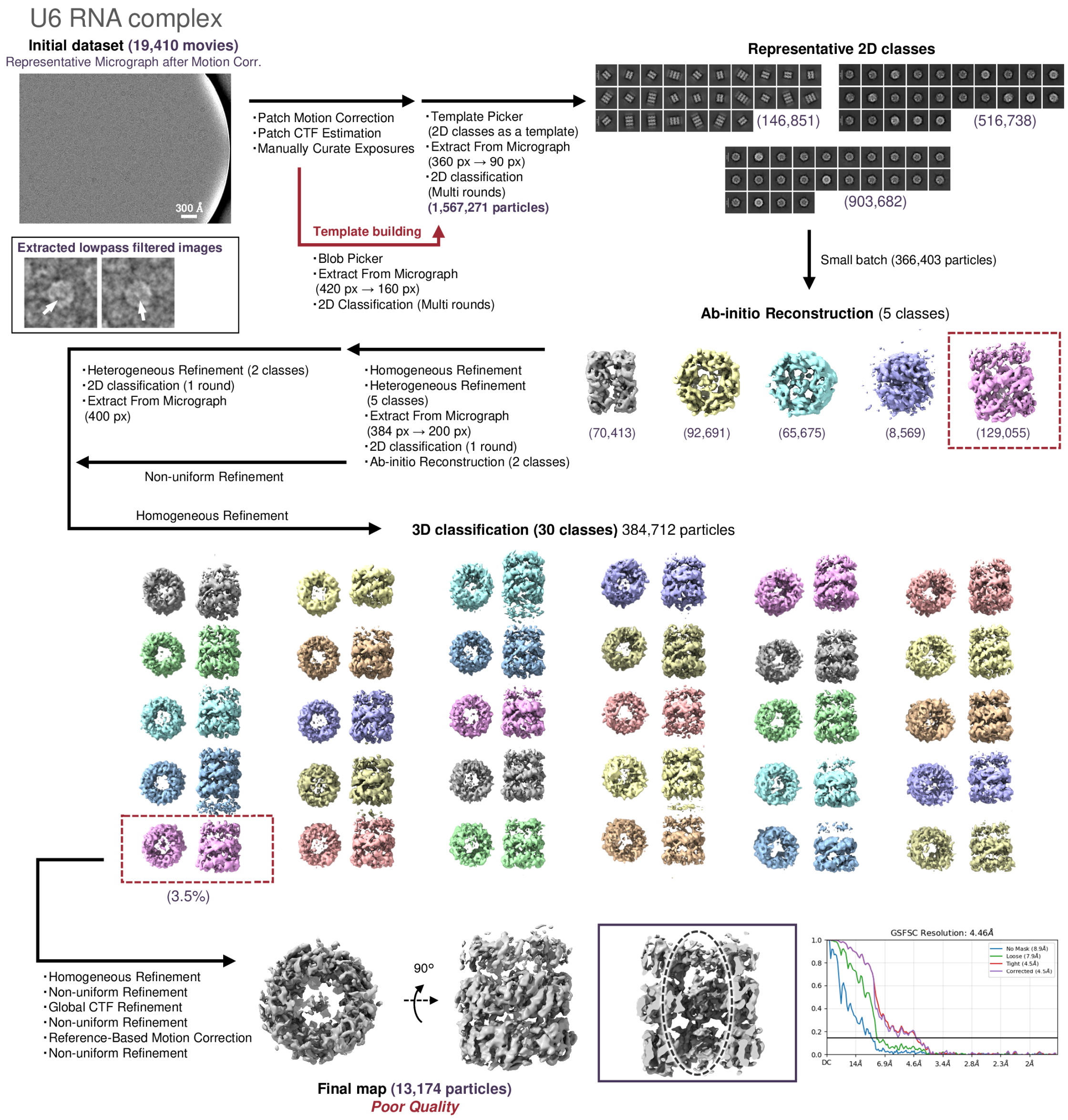

**Fig. S8 Single-particle cryo-EM processing workflow and reconstructions of U6 RNA complex.**

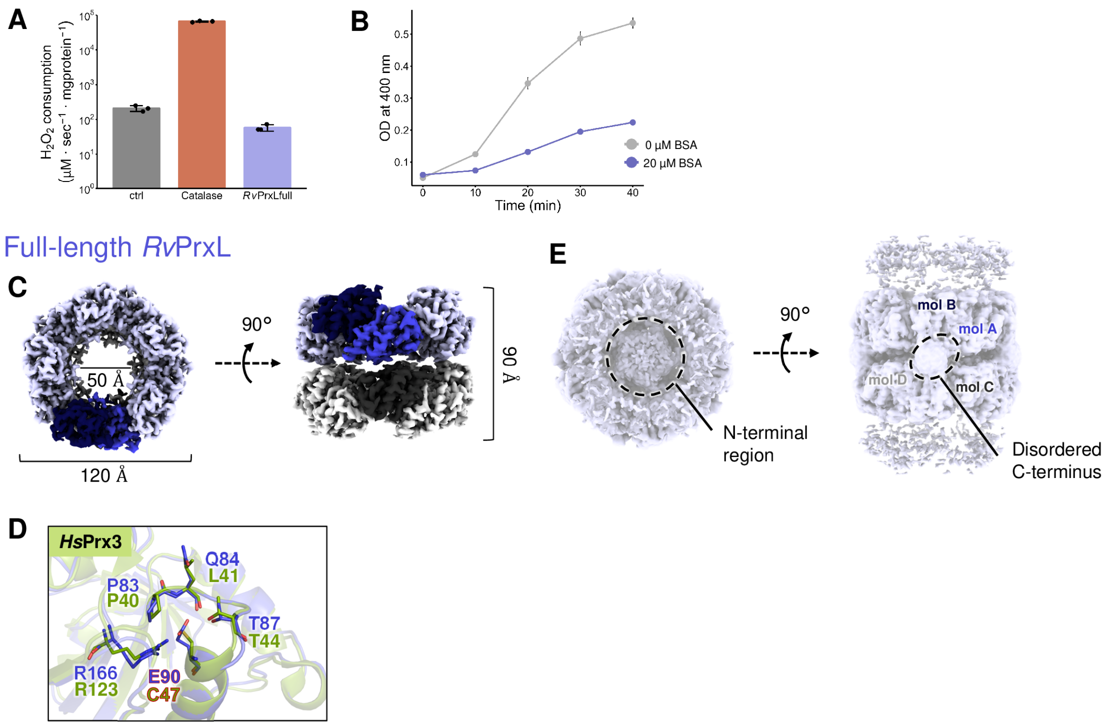

**Fig. S9 *Rv*PrxL activity assays and structural analysis of full-length *Rv*PrxL.**

(**A**) $H_2O_2$ consumption per minute of control, catalase, and *Rv*PrxL, measured by tracking the OD at 240 nm, which corresponds to the absorption wavelength of $H_2O_2$. (**B**) Assay control of holdase activity assay, employing bovine serum albumin (BSA). (**C**) Overall cryo-EM map of full-length *Rv*PrxL. (**D**) Structural comparison around E90 and $Cys^P$ residues, between *Rv*PrxL and *Hs*Prx3 (PDB ID: 5JCG). Mol A of *Rv*PrxL and the corresponding molecule of *Hs*Prx3 are used for comparison. (**E**) Low-threshold cryo-EM map of full-length *Rv*PrxL, revealing disordered N-terminal region and C-terminus. Residual densities above and below the 20-mer may be derived from the presence of higher order oligomeric assemblies such as 30-mer and 40-mer.

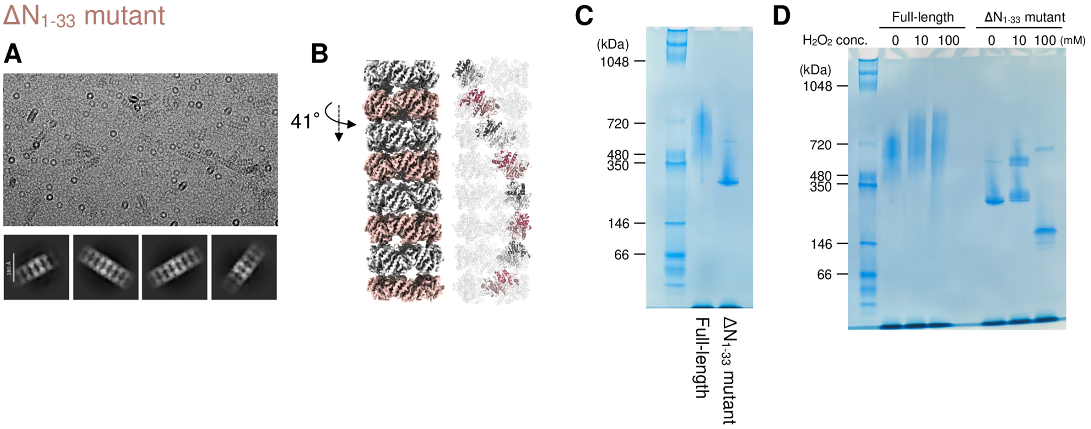

**Fig. S10 Structural analysis of $\Delta N_{1\text{–}33}$ mutant and Clear Native PAGE.**

(**A**) Representative micrograph and 2D classes, revealing the filamentous structure of $\Delta N_{1\text{–}33}$ mutant. (**B**) Overall map and ribbon representation model of $\Delta N_{1\text{–}33}$ mutant. 41° rotation between upper and lower 10-mers was observed, as found in the 20-mer of full-length *Rv*PrxL. (**C**) Clear Native PAGE (CN-PAGE) of full-length *Rv*PrxL and $\Delta N_{1\text{–}33}$ mutant, showing that full-length *Rv*PrxL exhibits a slightly higher molecular weight compared to $\Delta N_{1\text{–}33}$ mutant. (**D**) CN-PAGE of full-length *Rv*PrxL and $\Delta N_{1\text{–}33}$ mutant under the presence of 10 mM or 100 mM $H_2O_2$. Under the presence of 100 mM $H_2O_2$, $\Delta N_{1\text{–}33}$ mutant exhibited smaller molecular weight band, consistent with the result of cryo-EM analysis.

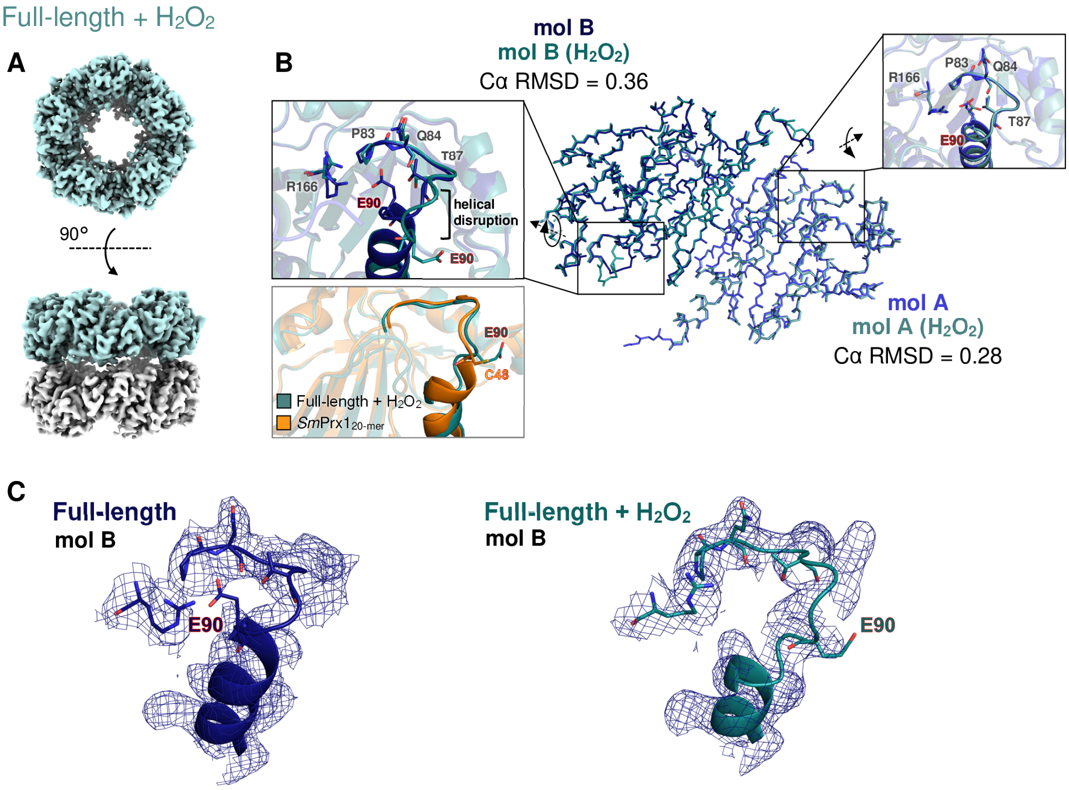

**Fig. S11 Structural analysis of $H_2O_2$-treated *Rv*PrxL.**

(**A**) Overall map of $H_2O_2$-treated *Rv*PrxL. (**B**) Structural comparison of dimer structures composed of mol A and mol B between *Rv*PrxL and $H_2O_2$-treated models. Comparisons around $Cys^P$ corresponding residues are indicated in the box. In the gray box, a structure around Cys48 (C48) in the 20-mer structure of *Sm*Prx1 (PDB ID:3ZVJ) is compared with a structure around Glu90 (E90) in $H_2O_2$-treated *Rv*PrxL. (**C**) Superposition of electropotential surface maps and structural models for the region surrounding Glu90 in structures with and without $H_2O_2$.

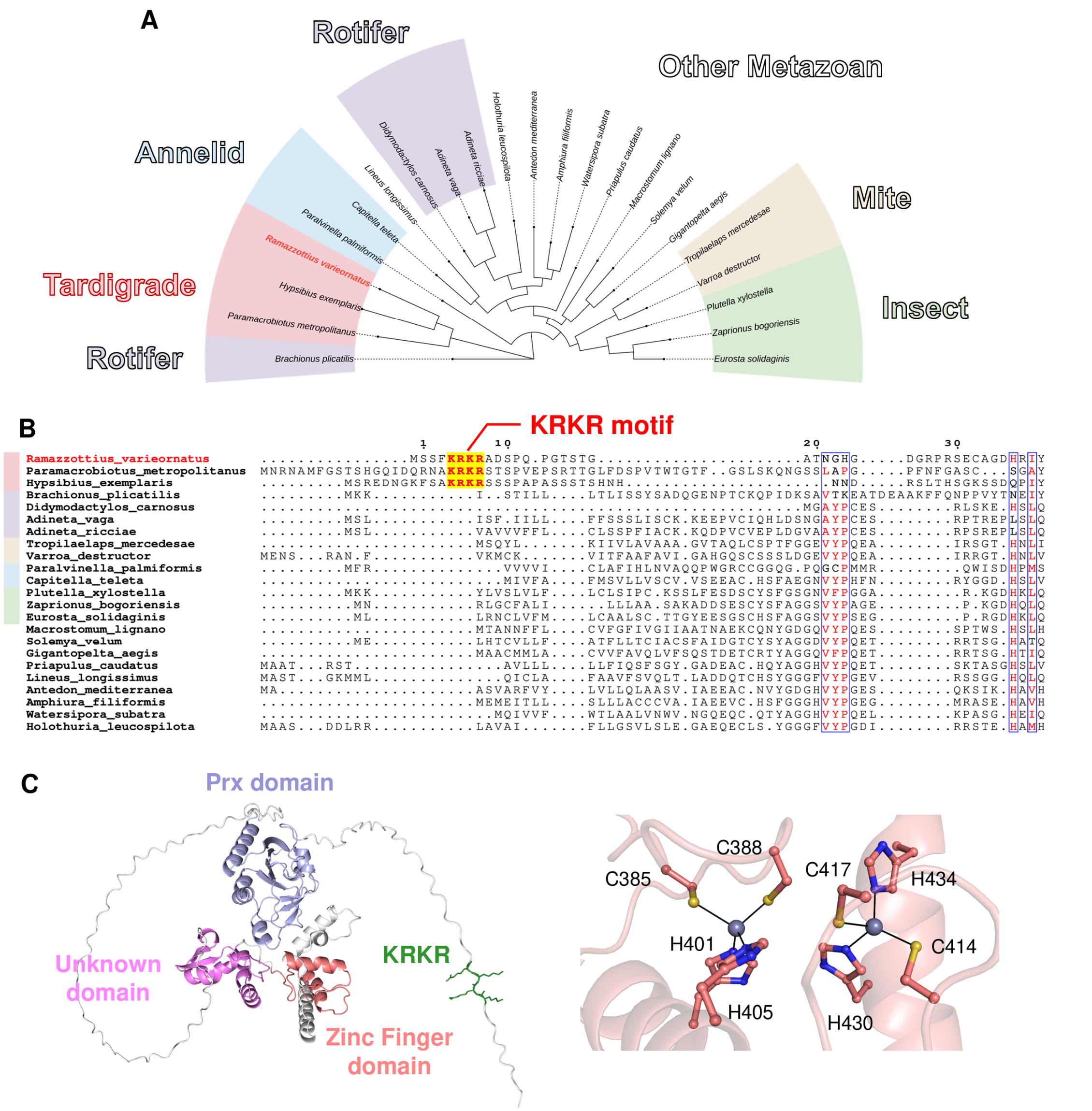

**Fig. S12 Tardigrade-specific KRKR motif revealed from comparative analysis of Prx homologs.** (**A**) Phylogenetic tree of high-scoring Prx homologs of *Rv*PrxL. Names of organisms are color-coded: pink, tardigrade; purple, rotifer; beige, mite; light blue, annelid; green, insect; and uncolored, other metazoans. (**B**) N-terminal sequence comparison and KRKR motif identification. N-terminal sequences were aligned using Clustal Omega. The conserved KRKR motif is present in all three homologous tardigrade Prx sequences. (**C**) Alphafold3-predicted structure of a zinc-finger domain containing Prx from *H. exemplaris* (Genbank: OQV14346.1). Zinc ions are shown as purple spheres in the right panel.

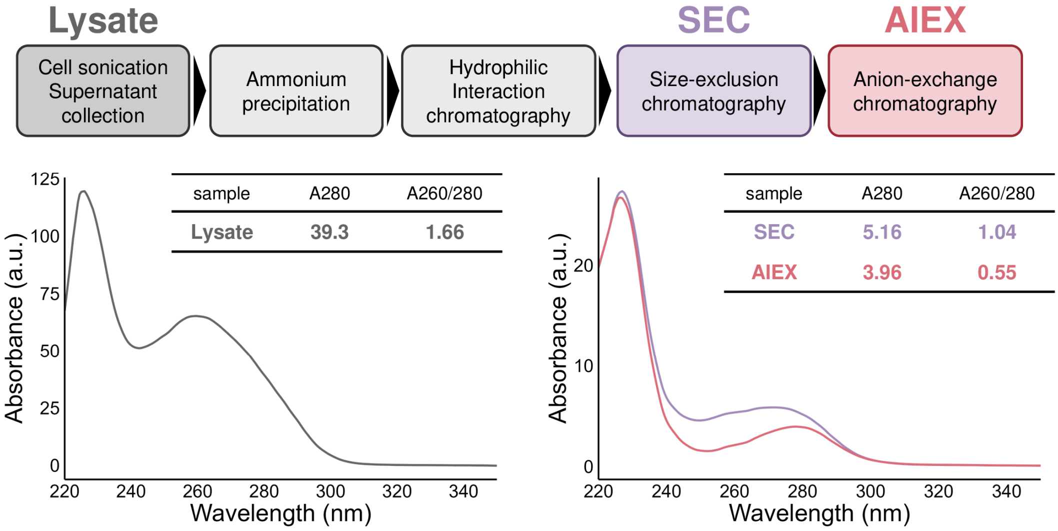

**Fig. S13 *Rv*PrxL purification process and UV-vis absorption spectroscopy.**

As purification proceeds, $A_{260/280}$ decreases, indicating the removal of nucleic acids contaminated in the sample.

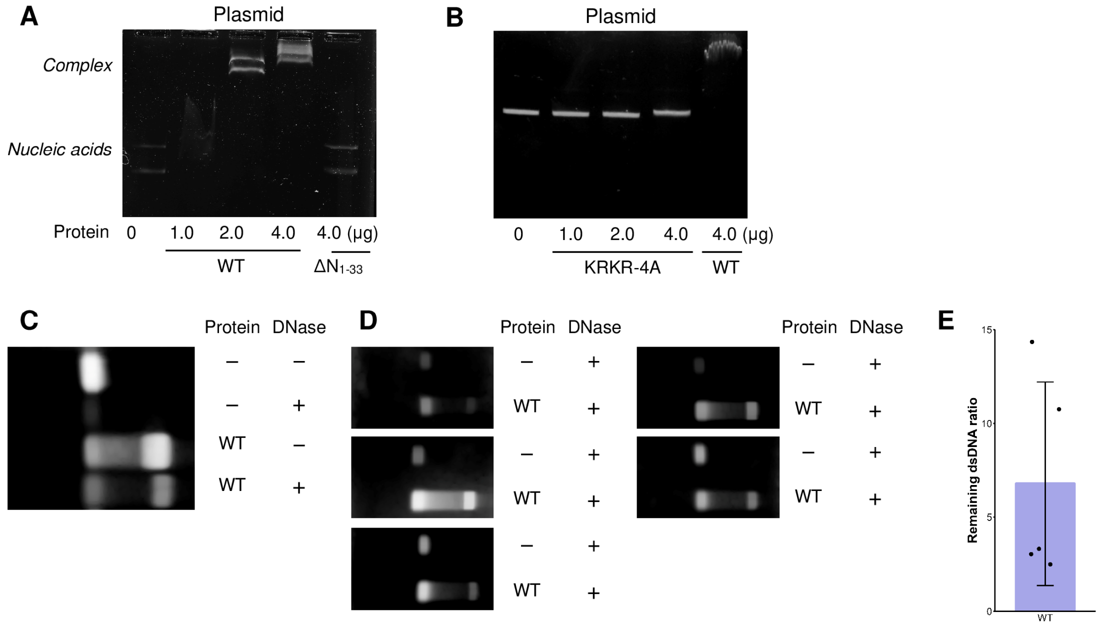

**Fig. S14 Mobility shift assay with plasmid and nuclease protection assay.**

(**A**) Mobility shift assay of full-length and $\Delta N_{1\text{–}33}$ mutant *Rv*PrxL with plasmid (pET-28a(+)). (**B**) Mobility shift assay of KRKR-4A mutant and full-length *Rv*PrxL with plasmid (pET-28a(+)). (**C**) DNase protection assay using dsDNA as a substrate. (**D**) Agarose gel images used to calculate the remaining dsDNA ratio. (**E**) Remaining dsDNA ratio relative to the control group. The remaining dsDNA intensity in the presence of *Rv*PrxL was, on average, 6.79 ± 4.85-fold higher than that in the control group.

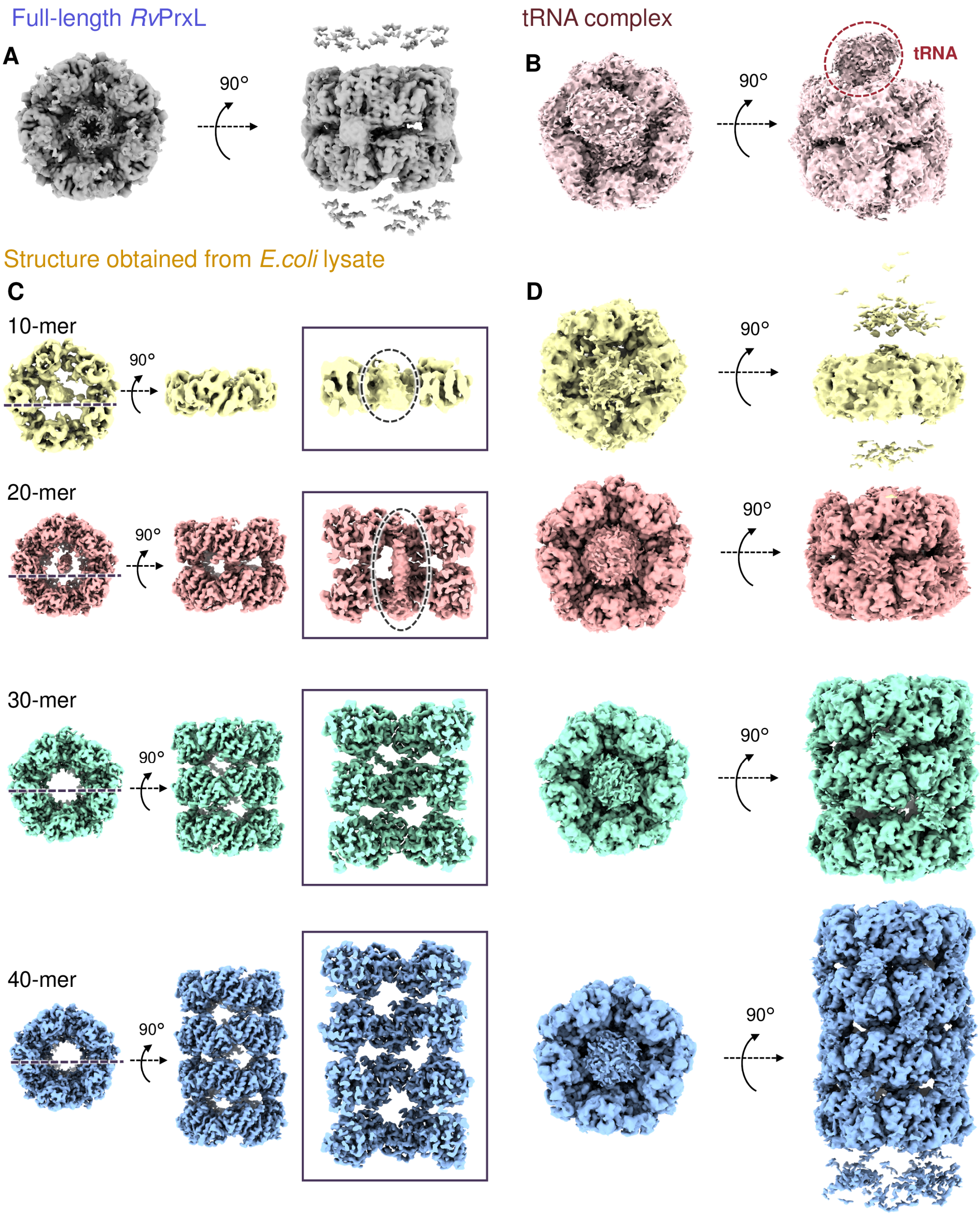

Full-length RvPrxL
A
90°
tRNA complex
B
90°
tRNA
Structure obtained from E.coli lysate
C
10-mer
90°
D
90°
20-mer
90°
90°
30-mer
90°
90°
40-mer
90°
90°

**Fig. S15 Low-threshold maps and cryo-EM maps of *Rv*PrxL structure obtained from *E. coli* lysate overexpressing *Rv*PrxL.**

(**A**) Low-threshold map of full-length *Rv*PrxL. (**B**) Low-threshold map of tRNA and *Rv*PrxL complex. (**C**) Overall map and vertical sectional view of the structure obtained from *E. coli* lysate. While 10-mer and 20-mer exhibit putative nucleic-acid structure, 30-mer and 40-mer have empty cavities, indicating that the nucleic acid only binds to the lower-oligomeric assembly. (**D**) Low-threshold map of the structure obtained from *E. coli* lysate overexpressing *Rv*PrxL.

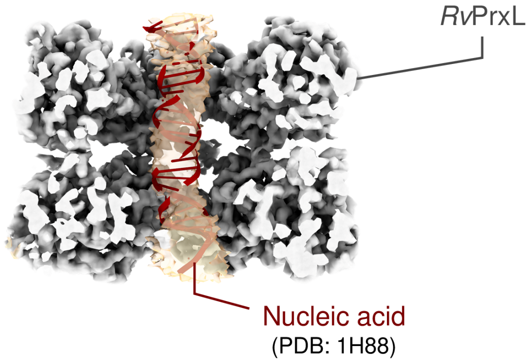

**Fig. S16 Superposition of a dsDNA model to putative nucleic acid density.**

**Supplementary Videos**

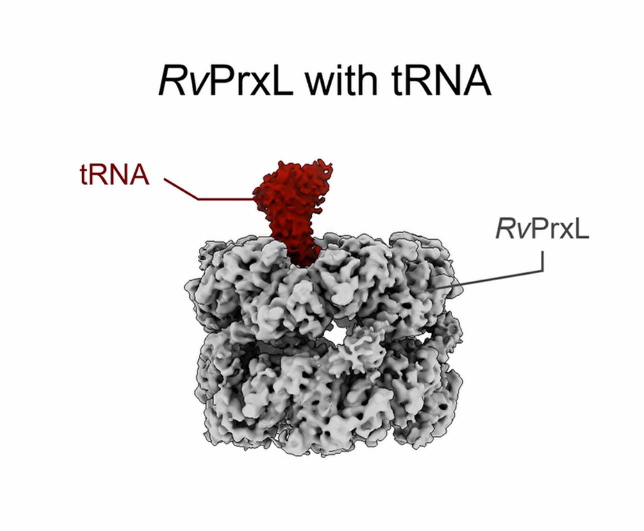

**Video S1 Cryo-EM map of the tRNA complex.**

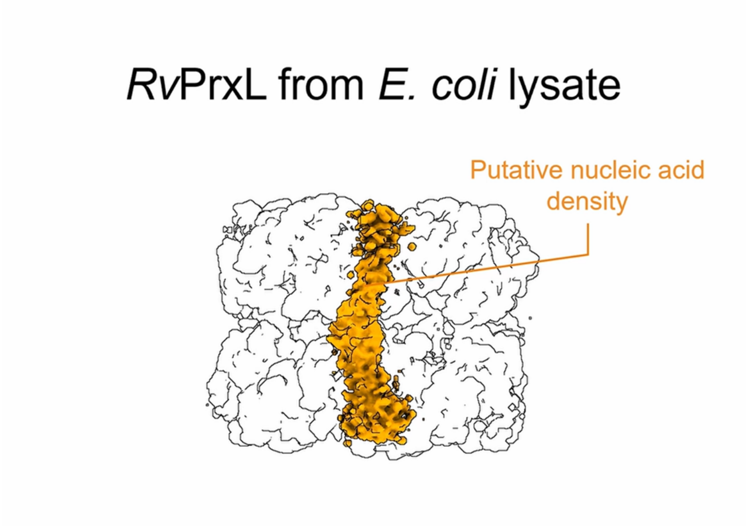

**Video S2 Cryo-EM map of a 20-mer with putative nucleic acid density.**
